# Trait anxiety drives premature disengagement despite intact opportunity-cost sensitivity

**DOI:** 10.64898/2026.06.19.733431

**Authors:** Peeusa Mitra, Garima Chauhan, Michael L. Platt, Arjun Ramakrishnan

## Abstract

Anxiety has been linked to difficulty sustaining engagement with ongoing tasks, even when continued engagement would yield greater rewards, yet the underlying mechanisms remain unclear. Here we examined how trait anxiety shapes sequential foraging decisions using a patch-leaving task grounded in the Marginal Value Theorem (MVT), a normative framework for explore–exploit decisions previously used to reveal altered computations in conditions such as problem gambling and attention-deficit hyperactivity disorder. Across two independent cohorts, participants adjusted patch residence times according to environmental opportunity costs, indicating preserved sensitivity to task structure irrespective of anxiety levels. Despite this, individuals with higher trait anxiety consistently left patches earlier and accrued fewer rewards. Drift diffusion modeling revealed that these deviations arose from reduced reward-driven evidence accumulation, rather than impaired environmental sensitivity or altered learning: trait anxiety reliably reduced the drift rate governing continued exploitation, providing a computational account of premature disengagement. This effect remained robust after accounting for pupil-linked arousal and baseline stress biomarkers, including salivary α-amylase and cortisol, which independently constrained decision dynamics. Together, these findings identify reduced reward-evidence accumulation as a core mechanism through which anxiety promotes premature disengagement from rewarding environments.

## Introduction

Anxiety and depression are highly prevalent and a major source of global disability [1]. Across these disorders, individuals often show reduced sustained engagement, impaired reward learning, and maladaptive decision-making under uncertainty [2,3]. One influential framework for studying such behavior is the exploration–exploitation trade-off, which captures the tension between persisting with a currently rewarding option and disengaging to seek potentially better alternatives [4,5]. Although explore–exploit paradigms have been widely applied to anxiety and depression, findings are mixed and difficult to interpret mechanistically [6,7]. In particular, it remains unclear whether anxiety alters how currently available rewards are used to guide continued engagement.

Patch-foraging tasks, grounded in optimal foraging theory, provide a principled framework for studying sequential engagement decisions [6]. Agents must decide whether to persist in a patch despite diminishing rewards or pay a cost to search for a new one. The Marginal Value Theorem (MVT) offers an elegant normative solution: leave when a patch’s instantaneous reward rate falls below the average reward rate of the environment [8–11]. MVT-consistent behavior has been observed across hundreds of species and contexts—from bees and birds to monkeys and humans—and across foraging for food, information, memory, strategies, and social resources [11–16], suggesting an evolutionarily conserved solution to deciding when to persist versus disengage. Because foraging requires sustained engagement with diminishing rewards under uncertainty, and recruits learning, executive control, valuation, and cognitive search, it provides a natural assay for the persistence-related disturbances characteristic of anxiety [17–21].

Stress is a major risk factor for affective disorders and biases foraging away from optimality, with acute and chronic stress promoting either excessive exploitation or premature switching depending on context [22,23]. Related disturbances appear in affective disorders—for example, increased exploration and reduced exploitation in major depression [24]—and anxiety and depression are increasingly understood as disorders of altered adaptation to uncertainty and reward [25,26]. A recent synthesis nonetheless emphasized substantial heterogeneity across tasks and outcomes, underscoring the need for mechanistic approaches linking stress-related traits to specific computational processes [7].

Trait anxiety offers a useful entry point for such an account [27]. Under the diathesis–stress model, stable individual differences in stress responsivity arise from interactions between genetic vulnerability and environmental stressors, particularly early in life [28]. Trait anxiety is marked by persistent worry, heightened sensitivity to uncertainty, and difficulty sustaining goal-directed engagement when continued investment yields diminishing or uncertain rewards [29]. Within a foraging framework, this could manifest as earlier patch-leaving driven not by impaired environmental knowledge but by altered accumulation and use of reward information.

At the neural systems level, stress shapes these decisions through neuromodulatory control of frontal and limbic circuits governing attention, salience evaluation, and behavioral persistence [30]. The sympathetic adreno-medullary system responds rapidly to stress, recruiting locus coeruleus–norepinephrine (LC–NE) signaling to mobilize arousal and reorient attention toward salient or uncertain events [31–33]. Through widespread cortical projections, norepinephrine regulates attentional gain, evidence accumulation, and exploration–exploitation trade-offs [34,35], and altered LC–NE function is repeatedly implicated in anxiety, including heightened tonic arousal, exaggerated responses to uncertainty, and reduced behavioral flexibility [36–38]. Stress also engages the slower hypothalamic–pituitary–adrenal (HPA) axis, whose end-product cortisol acts on prefrontal and limbic networks involved in executive control, salience processing, and reward-guided behavior, particularly under uncertainty or sustained demand [39–43]. Trait-anxious individuals frequently show dysregulation across both systems, including heightened cortisol reactivity and altered norepinephrine signaling [44,45].

These systems converge on a circuit linking the LC–NE system to the anterior cingulate cortex (ACC). The LC–NE system shifts between phasic firing, which promotes focused task engagement and selective processing of task-relevant information, and tonic firing, which favors disengagement and exploration [32,46,47]; under stress, elevated tonic activity heightens arousal and hypervigilance while reducing selectivity for currently available rewards, biasing behavior toward premature disengagement [48,49]. The dorsal ACC tracks reward rate, effort costs, and environmental utility during foraging and sequential choice [9,50–52], receives dense LC–NE input, and exerts reciprocal top-down control over the LC, allowing arousal states to be regulated by ongoing estimates of environmental value [53]. Because glucocorticoid receptors are densely expressed in medial prefrontal and cingulate cortex, cortisol-related HPA activity can further tune reward sensitivity, cognitive control, and decision thresholds within this circuit [43,54]. Together, these interacting ACC–LC–cortisol pathways provide a plausible substrate through which anxiety and stress may reduce reward-guided persistence and promote premature patch leaving.

Patch-leaving can be framed computationally as sequential evidence accumulation, in which agents integrate the declining value of the current option against the opportunity cost of searching, disengaging when accumulated evidence crosses a decision threshold [9,55,56]. This process is sensitive to stress and arousal: neuromodulatory systems can alter either the rate of reward-evidence accumulation or the disengagement threshold, biasing decisions away from normative solutions [41,43,46,49]. Pupillometry provides a noninvasive window onto these dynamics, with tonic pupil size indexing baseline arousal and disengagement readiness and phasic dilation reflecting transient reward and attentional responses; pupil fluctuations also track latent decision variables such as uncertainty and exploration–exploitation state [57–61].

Here we combined behavioral, computational, and physiological approaches across two independent cohorts (India and the United States) who completed an MVT-based patch-foraging task alongside trait-anxiety assessment (STAI-Y2). In a subset, we additionally measured pupil dynamics and baseline salivary biomarkers—α-amylase (sAA; LC–NE/sympathetic arousal) and cortisol (HPA-axis activity). We hypothesized that trait anxiety would bias foraging not by disrupting sensitivity to opportunity costs, but by altering how reward information is accumulated into decisions to persist or disengage—specifically, that anxiety-related stress physiology would reduce effective reward-evidence accumulation during exploitation, promoting premature disengagement despite preserved opportunity-cost sensitivity.

## Materials and Methods

### Participants

Two independent cohorts were studied. The U.S. cohort comprised 137 healthy adults (40 male, 97 female; age 19–45 y) tested at the Wharton Behavioral Laboratory, University of Pennsylvania; the study was approved by the University of Pennsylvania Institutional Review Board and all participants provided informed consent. The Indian cohort comprised 55 healthy adults (37 male, 18 female; age 18–32 y) recruited from the student population at the Indian Institute of Technology (IIT) Kanpur; the study was approved by the Institute Human Ethics Committee of IIT Kanpur (IITK/IEC/2020-21/II/10) and all participants provided written informed consent. All participants were in good health and reported no medications affecting cognitive performance. U.S. sessions lasted ∼20 min (sequential patch-foraging task, ∼180–200 trials). Indian sessions lasted ∼1 h and additionally included electroencephalography (EEG), eye-tracking, and saliva collection; to minimize circadian variation, all Indian sessions were run between 12:00 and 19:00. Quality-control exclusion criteria and final analytic samples are detailed in Supplementary Methods (final behavioral samples: U.S. n = 79, India n = 55; hierarchical sequential sampling models: U.S. n = 123, India n = 55).

### Foraging task

Participants performed a computer-based patch-foraging task (Fig. 1A–C) adapted from established paradigms [6,9,70]. After written instructions and a practice block (with different parameters), participants were told their monetary bonus was proportional to total berries collected. On each trial they chose to harvest the current patch or leave to search for a new one, implemented through a two-step fixation procedure that isolated a pre-decisional (tonic) period: an initial click followed by a 1-s wait, then a confirmatory click once the fixation cue turned dark green. Harvesting yielded a reward; leaving initiated a travel delay (downward screen scrolling) that operationalized environmental opportunity cost. Travel times defined “rich” (short) and “poor” (long) environments and were learned through experience rather than instructed.

**Figure 1.**
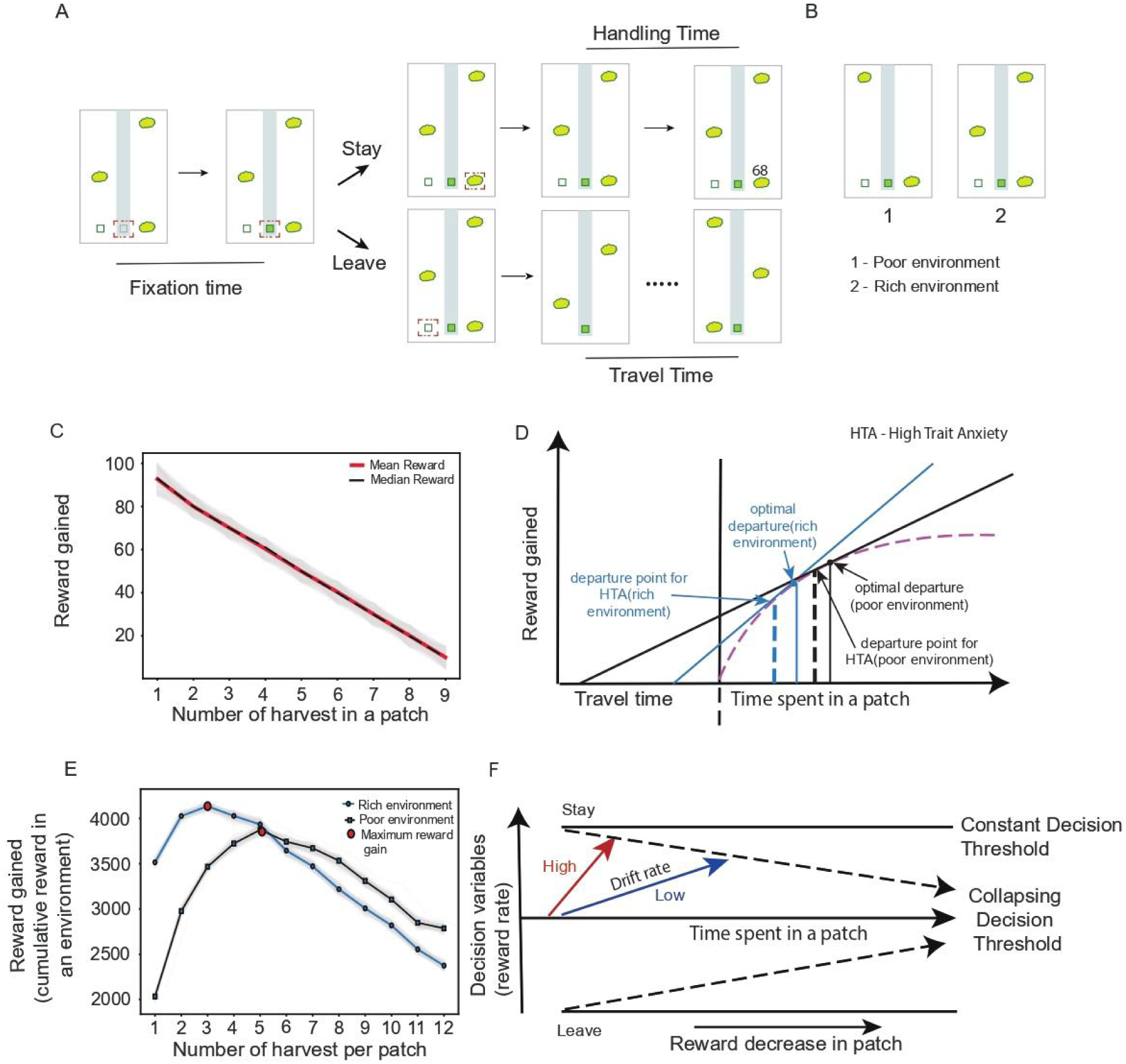
Patch-foraging task, reward structure, and computational framework. (A) Schematic of the sequential patch-foraging task. Each trial began with a fixation requirement, after which participants chose to either harvest from the current patch or leave. Harvesting yielded rewards that decayed with successive selections, whereas leaving initiated a travel delay before entering a new patch. Each block lasted 5 minutes. (B) Environmental manipulation of opportunity cost. “Rich” and “poor” environments were defined by shorter (3 s) and longer (10 s) travel times between patches, respectively. (C) Reward decay within a patch as a function of harvest number. The solid line represents mean reward and the dashed line represents median reward across trials. Shaded regions indicate variability in reward depletion due to stochasticity. (D) Depiction of the Marginal Value Theorem and the optimal time to leave a patch as a function of between-patch travel duration. Increased travel duration calls for longer stays. Bold perpendiculars indicate the early departure of high-trait-anxious individuals. (E) Total reward accumulation as a function of harvest number for rich (3 s) and poor (10 s) travel-time environments. Curves illustrate environment-specific optimal foraging policies, with peaks (red dots) indicating the harvest number that maximizes reward accumulation. (F) Computational framework. Schematic of a drift diffusion model, with drift rate reflecting reward-dependent evidence accumulation to stay or leave. Drift rate decreases as available reward declines. The collapsing decision threshold depicts the decrease in threshold with reward decrement.

For the U.S. cohort, the initial patch reward was r0 = 7 + N(0, 0.25), depleting as ri = ri−1 − 0.5 + N(0, 0.25), bounded at zero; two 4-min blocks used 5-s (short) and 20-s (long) travel times. For the Indian cohort, initial rewards were sampled uniformly between 85 and 100 and depleted by a fixed decrement (ε = 10) with added stochasticity, Ri = rand(R_{i−1}upper − ε, R_{i−1}lower − ε), bounded at zero; two 5-min blocks used 3-s (short) and 10-s (long) travel times. The task was implemented in PsychoPy on a 27-inch monitor (1920 × 1080, 60 Hz). Full reward-structure and task details are provided in Supplementary Methods.

### Self-report, pupillometry, and salivary biomarkers

Participants completed the Spielberger State–Trait Anxiety Inventory (STAI; 20 items per subscale, 4-point Likert). Trait anxiety (STAI-Y2) was assessed once at baseline and used as the individual-difference index; state anxiety (STAI-Y1) was assessed before and after the task. Pupil diameter (left eye) was recorded at 200 Hz (Pupil Core, Pupil Labs) and synchronized to behavioral/EEG markers via Lab Streaming Layer. After blink removal, interpolation, and smoothing (PyPLR), tonic pupil size was defined as mean diameter from −750 to −250 ms before the center-cue click, and phasic responses from 1000–1250 ms after the decision. Saliva was collected at baseline (T1) and post-task (T2) using Salivette kits and stored at −80 °C; cortisol and α-amylase were quantified by ELISA in duplicate, with baseline (T1) values indexing HPA-axis and noradrenergic tone, respectively. Full acquisition, preprocessing, epoch-extraction, and assay details are provided in Supplementary Methods.

### Statistical analysis

Analyses used R 4.4.1. Bayesian regression models (brms) tested how patch-stay duration and patch-wise cumulative reward varied with travel time × trait anxiety, and how trial-level tonic and phasic pupil measures varied with reward-related variables and trait anxiety (random intercepts for participants). Mediation analyses (mediation, lavaan; quasi-Bayesian Monte Carlo, 500 simulations) tested whether pupil-linked arousal accounted for the trait anxiety–reward relationship and were interpreted as statistical mediation without implying causality. To recover latent decision processes, hierarchical sequential sampling models (HSSM v0.2.11; Bambi, NumPyro, PyMC, ArviZ) jointly modeled stay–leave choices and response times (RTs), estimating drift rate (v; reward-evidence accumulation), boundary separation (a; decision caution), and starting point (z). Trial-wise predictors (reward, travel time, phasic pupil, trait anxiety, and baseline sAA and cortisol) were entered on v and a across models of increasing complexity. To benchmark optimality, observed behavior was compared with MVT-derived optimal harvest numbers, and patch-wise reward deviation was computed as the difference between actual and model-optimal cumulative reward. Full model specifications, equations, and priors are provided in Supplementary Methods.

## Results

### Trait anxiety predicts premature patch leaving and reduced reward accumulation

We compared patch-leaving behavior with MVT predictions, which specify how long to persist in a patch given the opportunity cost of travel between patches; MVT reduces a complex sequential problem to local reward rate and global opportunity cost (Fig. 1D,E). In the Indian cohort (n = 55), participants stayed longer in the poor environment, where travel time,and thus opportunity cost was higher: mean stay was 12.0 ± 5.42 s in the rich environment (travel = 3 s; MVT-optimal = 11.96 s) and 17.19 ± 5.19 s in the poor environment (travel = 10 s; MVT-optimal = 21.67 s). This replicated in the U.S. cohort (n = 137): 30.89 ± 15.30 s in the rich environment (travel = 5 s; optimal = 23.44 s) and 46.88 ± 20.75 s in the poor environment (travel = 20 s; optimal = 38.57 s). Longer absolute durations in the U.S. reflected longer trial durations in that implementation (mean trial duration: India 3.81 ± 1.42 s; U.S. 5.12 ± 2.04 s).

Despite this preserved sensitivity, patch-leaving varied substantially across individuals (Indian IQRs: 9.05–15.34 s rich, 13.99–20.25 s poor; U.S. IQRs: 20.55–39.43 s rich, 33.92–58.21 s poor).

To test whether this variability tracked trait anxiety, particularly the tendency toward premature patch leaving, we fit Bayesian regressions of patch-wise stay duration on travel time and trait anxiety. Longer travel times reliably increased stay duration in both cohorts (India(n = 55): β = 0.38, 95% CI [0.35, 0.41]; U.S.(n = 79): β = 0.36 [0.32, 0.40]). Critically, higher trait anxiety predicted earlier disengagement in both (India(n = 55): β = −0.08 [−0.11, −0.05]; U.S.(n=79): β = −0.09 [−0.13, −0.06]; Supplementary Tables S1–S2). Trait anxiety did not interact with travel time (Supplementary Fig. S1), indicating preserved sensitivity to opportunity costs across anxiety levels, and including anxiety improved model fit (ΔELPD = 22.8, SE = 7.9).

These behavioral differences translated into reward outcomes. Reward harvest increased with travel time (India(n =55): β = 0.35 [0.30, 0.40]; U.S.(n = 79): β = 0.21 [0.18, 0.25]), while higher trait anxiety was associated with reduced per-patch reward in both cohorts (India: β = −0.11 [−0.16, −0.06]; U.S.: β = −0.07 [−0.11, −0.04]; Supplementary Tables S3–S4, Supplementary Fig. S2). Thus, anxious individuals remained sensitive to opportunity costs yet disengaged earlier, accruing less reward.

### Rewards, trait anxiety, and stress neuromodulators shape pupil dynamics

To probe physiological processes underlying these differences, we examined pupil dynamics as a readout of arousal and neuromodulatory state during foraging. We quantified how reward variables shaped pupil size across three timescales, local reward on individual harvests (trial level), cumulative reward within a patch (patch level), and global context (travel time, average reward rate), separately for tonic (1 s pre-decision) and phasic (post-decision) pupil (Fig. 2A).

**Figure 2.**
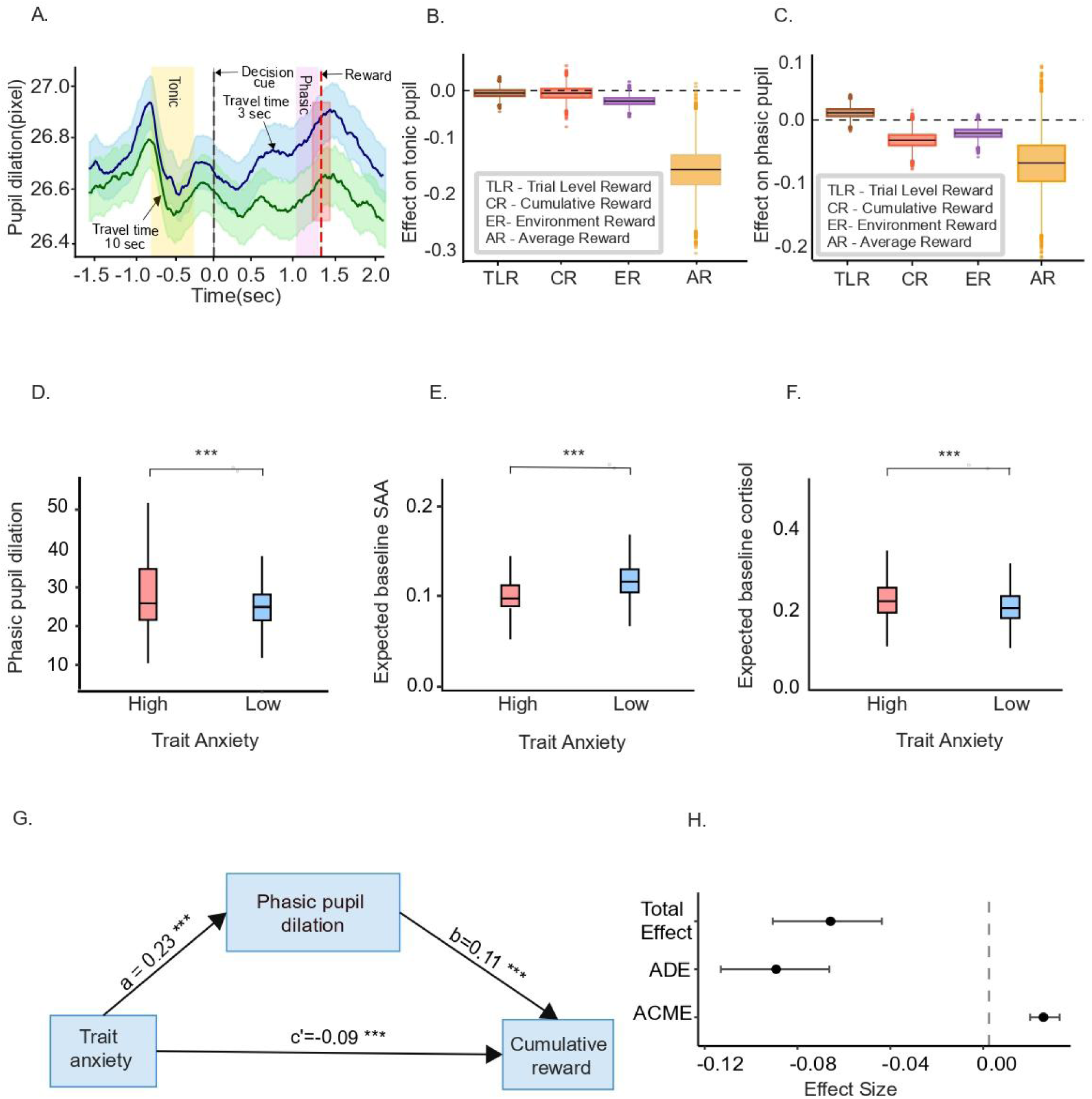
Pupil-linked arousal, neuromodulatory state, and their relationship to trait anxiety and reward accumulation. (A) Trial structure and pupil dynamics. Schematic illustrating tonic (pre-decisional) and phasic (post-decisional) pupil responses aligned to decision events. (B–C) Effects of reward-related variables on pupil size across timescales. Regression coefficients (notched box plots) show the influence of trial-wise reward (brown), cumulative reward within a patch (peach), environmental reward rate (violet), and average reward (yellow) on tonic (B) and phasic (C) pupil dilation. Notches indicate 95% confidence intervals of the median; whiskers represent the full range. Negative coefficients (below the dashed line) indicate reduced pupil size with increasing reward signals. (D–F) Trait anxiety and physiological markers. Group comparisons (high vs low trait anxiety) for phasic pupil dilation (D), baseline salivary α-amylase (sAA; E), and baseline cortisol (F). Higher-trait-anxiety individuals exhibit increased phasic pupil responses and altered neuromodulatory profiles. Cortisol differences were derived from Bayesian regression models (baseline cortisol ∼ trait anxiety). Notched box plots show median and 95% confidence intervals. Asterisks (***) denote significant effects; “ns” indicates non-significance (see Supplementary Information). (G) Mediation model. Path diagram illustrating the indirect relationship between trait anxiety and cumulative reward via phasic pupil dilation. Solid arrows indicate significant paths; standardized coefficients are shown. The direct effect of trait anxiety is denoted by c′. (H) Mediation effects. Point estimates and 95% confidence intervals for the average causal mediation effect (ACME), average direct effect (ADE), and total effect. The vertical dashed line indicates zero effect.

At the trial level, reward robustly increased phasic dilation (β = 0.02 [0.01, 0.04]), strongest for high-value rewards early in a patch (Fig. 2C), but did not modulate tonic size (Fig. 2B). At the patch level, cumulative reward was associated with reduced phasic dilation (β = −0.02 [−0.04, 0.00]; Fig. 2C), consistent with diminishing marginal utility as exploitation continued, again without affecting tonic size. At the environmental level, longer travel times reduced both phasic (β = −0.01 [−0.03, −0.00]) and tonic pupil (β = −0.02 [−0.03, −0.00]), and higher average reward rates robustly reduced tonic size (β = −0.15 [−0.23, −0.06]) without affecting phasic dilation (Fig. 2B,C). Thus phasic responses tracked local reward events whereas tonic size tracked global statistics; a temporal dissociation matching the timescales over which MVT-relevant variables evolve.

These signals were systematically shaped by anxiety and stress physiology. High-trait-anxiety individuals showed greater phasic dilation (Fig. 2D), and trait anxiety predicted both tonic (β = 0.69 [0.61, 0.76]) and phasic pupil (β = 0.72 [0.65, 0.79]; Supplementary Table S5A, Supplementary Fig. S3). With baseline sAA in the model, trait anxiety remained a robust predictor of phasic dilation (β = 0.96 [0.88, 1.05]) and sAA itself amplified phasic responses (β = 0.75 [0.59, 0.91];Supplementary Image S6; Supplementary Table S5B), whereas baseline cortisol was associated with reduced phasic dilation (β = −0.26 [−0.32, −0.21]; Supplementary Image S7; Supplementary Table S5C) indicating partially dissociable LC–NE and HPA contributions. Finally, mediation analysis showed that trait anxiety increased phasic dilation (a-path: 0.23 ± 0.01, p ≤ 0.001), which in turn predicted cumulative reward (b-path: 0.10 ± 0.01, p ≤ 0.001), with a significant average causal mediation effect (ACME β = 0.02 [0.01, 0.02]; Fig. 2G,H; Supplementary Table S6). Comparable effects held for tonic pupil (Supplementary Table S7, Supplementary Fig. S4), supporting pupil-linked arousal as a statistically informative intermediary associated with the relationship between trait anxiety to foraging.

### Trait anxiety alters the computational dynamics of patch leaving

While the preceding analyses established that trait anxiety was associated with premature patch leaving, identifying the underlying computational mechanism requires a process-level account linking MVT-based foraging behavior to internal decision dynamics. We therefore fit drift diffusion models (DDMs) to jointly model stay–leave choices and response times (RTs)[9,55,56]. Within the context of patch foraging, the DDM provides a mechanistic implementation of the MVT by formalizing patch leaving as a sequential evidence-accumulation process in which reward information supporting continued exploitation is accumulated over time until a decision criterion favoring disengagement is reached [9,62]. In this framework, Drift rate (v) indexes reward-evidence accumulation favoring continued exploitation whereas boundary separation (a) captures the amount of evidence required before a choice is made (Fig. 1F).

Because choices alone reveal only whether a participant stayed or left, they provide limited insight into the latent process generating those decisions. Response times, in contrast, contain information about the dynamics of evidence accumulation leading up to each choice. Furthermore, response times varied substantially across trials and individuals and were associated with patch-stay duration (Supplementary Results S9.a, Indian cohort; S9.b, U.S. cohort).

We first examined whether the latent decision variables recovered by the model reflected key determinants of optimal foraging predicted by the MVT. In the Indian cohort (n = 55), drift rate increased with marginal reward size (B = 1.798 [1.727, 1.870], p < 0.001) and with longer travel times (B = 0.075 [0.054, 0.094], p < 0.001), as predicted by MVT (a poorer environment raises the relative value of the current patch). Critically, trait anxiety significantly reduced drift rate (B = −0.085 [−0.149, −0.021], p < 0.001; Fig. 3A; Supplementary Table S14), indicating weaker reward-evidence accumulation. The U.S. cohort (n = 123) showed the same pattern: drift increased with marginal reward (B = 1.484 [1.458, 1.509], p < 0.001) and travel time (B = 0.024 [0.021, 0.027], p < 0.001), and trait anxiety significantly reduced drift (B = −0.033 [−0.054, −0.010], p < 0.001; Fig. 3C; Supplementary Table S19).

**Figure 3.**
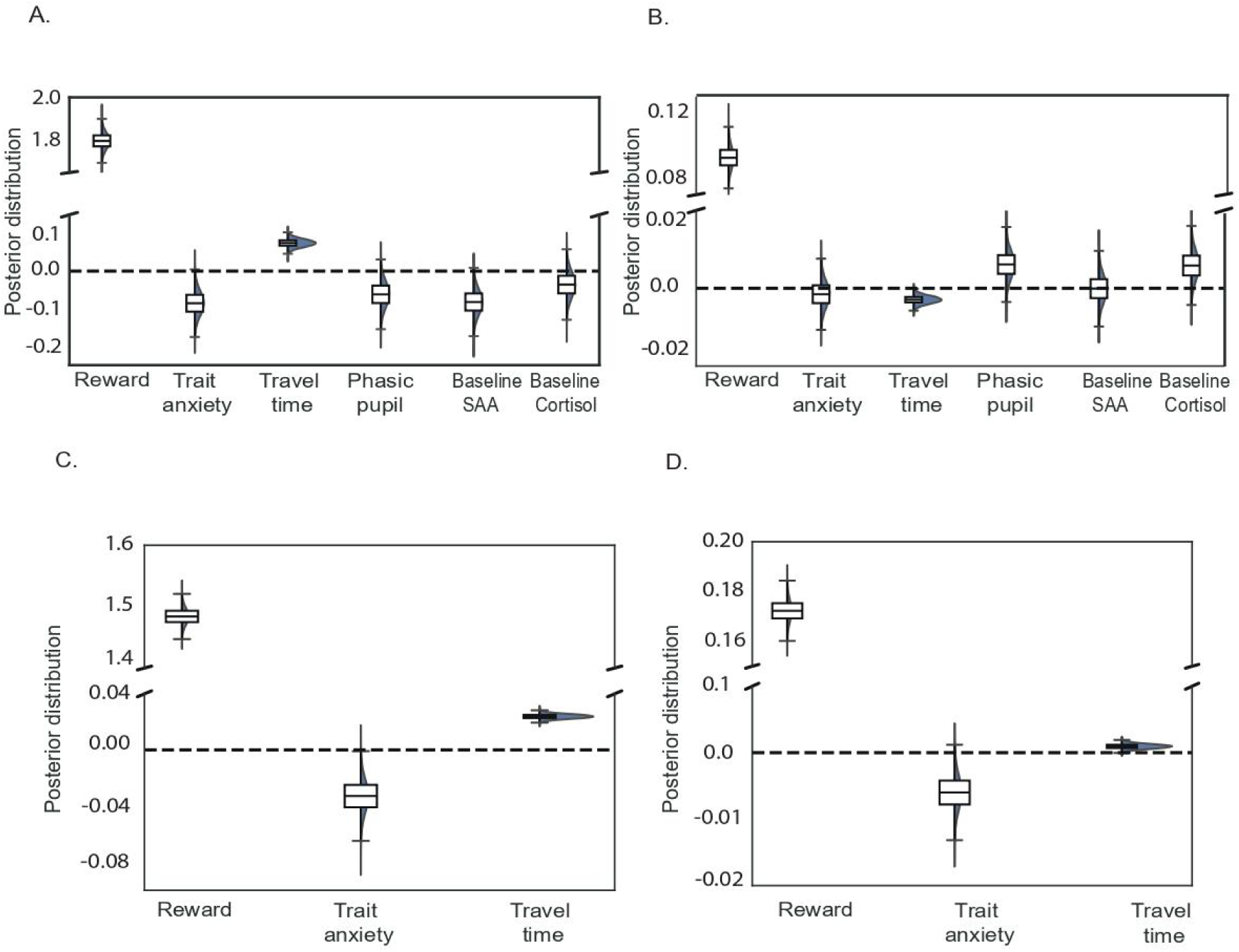
Latent decision dynamics underlying patch leaving: effects of reward, arousal, and neuromodulatory state. Posterior estimates from hierarchical sequential sampling models (HSSM) illustrate how task variables, pupil-linked arousal, trait anxiety, and neuromodulatory markers modulate latent decision parameters. (A, C) Effects on drift rate (v). Trial-wise reward, environmental opportunity cost (travel time), phasic pupil dilation, trait anxiety, and baseline salivary markers (α-amylase and cortisol) modulate the rate of reward-related evidence accumulation. (B, D) Effects on decision boundary (a). The same predictors influence decision thresholds, reflecting changes in decision caution during patch-leaving choices. Violin plots depict posterior distributions of regression coefficients, with central dashed lines indicating posterior means and distribution width reflecting uncertainty. The horizontal dashed line at zero indicates no effect. Colors are used for visualization only and do not encode specific conditions.

Threshold effects were less consistent. Higher local reward raised thresholds in both cohorts (India: B = 0.093 [0.079, 0.106]; U.S.: B = 0.172 [0.164, 0.181]), but travel time exerted small opposing effects (India: B = −0.004 [−0.006, −0.001]; U.S.: B = 0.001 [0.000, 0.002]), and the trait-anxiety effect did not replicate (India: B = −0.002 [−0.010, 0.006], p = 0.61; U.S.: B = −0.006 [−0.011, −0.001]). Thus, unlike drift rate, decision thresholds were not a generalizable signature of anxious foraging.

Adding pupil measures improved model fit (pLOO = 10.39; Supplementary Table S16): larger phasic responses predicted reduced drift (B = −0.062 [−0.126, 0.002] tending to significance), with similar effects for tonic pupil (Supplementary Table S14) and smaller, less consistent threshold effects. Adding baseline stress biomarkers(salivary cortisol and sAA improved fit further (pLOO = 14.30): higher baseline sAA (B = −0.082 [−0.147, −0.021]) and cortisol (B = −0.036 [−0.102, 0.029], tending to significant) both reduced drift (Fig. 3A), mirroring the anxiety effect, again with less consistent threshold effects (Fig. 3B; Supplementary Table S14).

Together, across two independent cohorts higher trait anxiety consistently reduced drift rate, diminished accumulation of reward evidence during exploitation; while larger pupil responses and elevated baseline cortisol and sAA showed convergent associations with reduced drift, linking stress-related physiological arousal to reduced reward-driven persistence.

Full posterior summaries for all drift-rate and boundary-separation effects, across cohorts and model variants, are provided in Supplementary Tables S14–S19.

## Discussion

We examined how trait anxiety shapes sequential decision-making by combining a normative ecological framework with computational modeling and physiological measures of arousal and stress. Trait anxiety biased decisions by weakening the influence of reward information on evidence accumulation during ongoing engagement, producing premature disengagement despite preserved sensitivity to opportunity costs. Behavior scaled with opportunity costs as predicted by the MVT [62], and even high-trait-anxiety individuals adjusted patch residence to travel time—indicating intact encoding of environmental structure—yet they left earlier and accrued less reward [63].

Computational modeling localized this effect to reduced drift rates governing reward-evidence accumulation during exploitation [56]. Lower drift indicates weaker integration of reward information supporting continued engagement, providing a direct account of premature leaving [64]. Although reduced drift rate could in principle arise from altered subjective valuation, the preservation of opportunity-cost sensitivity across anxiety levels suggests that anxious individuals retain accurate representations of environmental structure, with the primary difference emerging in the influence of experienced rewards on persistence decisions. Our findings do not exclude altered subjective valuation in anxiety [65] but suggest such effects manifest in how reward information is accumulated and used to sustain engagement under uncertainty [26,66]. Diminished reward-driven accumulation emerged as the primary computational signature across both cohorts [67,68], and it remained robust after accounting for pupil dynamics and baseline biomarkers, indicating that trait anxiety contributes independently to altered evidence accumulation rather than being fully mediated by physiological arousal.

Physiological measures provided convergent but partially independent constraints. Phasic pupil responses tracked experienced rewards, whereas tonic pupil size tracked broader context (travel time, average reward rate), consistent with pupil-linked arousal indexing neuromodulatory control of uncertainty and decision policy [58,60,69,70]. Larger pupil responses predicted reduced drift, aligning with literature linking pupil dilation to LC–NE activity and exploration–exploitation regulation [46,49,71] and to subjective value, reward volatility, and effort costs [72,73]. Baseline biomarkers revealed dissociable influences: sAA amplified phasic responses under higher anxiety, whereas cortisol elevated tonic arousal while dampening phasic reactivity—mirroring prior links between cortisol and sympathetic arousal and stress-related changes in control and attention [43,74,75], and supporting models in which anxiety reflects dysregulated attentional and control systems rather than disturbed reward valuation alone [29,76].

These results fit current models of stress-related neuromodulation in frontal control circuits, in which the ACC monitors reward rate and regulates persistence while the LC–NE system shifts between phasic (engaged) and tonic (disengaged/exploratory) modes; under stress, elevated tonic LC–NE activity reduces reward-driven evidence accumulation [9,46,48–51,53]. Within this framework, the reduced drift rates of anxious individuals may reflect weakened reward signaling within ACC-mediated persistence circuits. Sequential foraging is well suited to reveal such influences because local stay–leave decisions are embedded in broader contexts, permitting dissociation of moment-to-moment reward processing from longer-timescale engagement—which may help explain why anxiety-related arousal biases persistence without abolishing sensitivity to contingencies [77].

Several limitations apply. Mediation and moderation analyses identify pupil dynamics as statistically informative intermediaries but cannot establish causality, and pupil size is an indirect proxy for neuromodulatory activity; direct neural recordings or causal manipulations of stress systems would strengthen mechanistic inference. Nevertheless, the convergence of behavioral, computational, physiological, and hormonal measures provides a robust multi-level account.

In summary, anxiety-related deviations from optimal foraging arise not from impaired sensitivity to opportunity costs but from stress-related shifts in internal control states that reduced reward-evidence accumulation during ongoing engagement. By integrating mathematical ecology, computational modeling, pupillometry, and stress physiology, this work identifies premature disengagement from rewarding environments as a core feature of anxious decision-making and generates clear translational predictions: interventions targeting tonic arousal regulation or enhancing persistence during reward harvesting may help restore engagement with objectively rewarding activities [78].

## Data Availability Statement

The data that support the findings of this study, together with the analysis code, are available from the below link. [https://osf.io/4kc9j/overview?view_only=015d5d5b82be48fca911606c7a4763e3]

## Acknowledgments

We thank Santosh Misra for access to equipment used in salivary biomarker analyses. We are grateful to Leafy Behera, Pratyusha Chakraborty, and Khyathi Vagolu for assistance with data collection, pupil analysis pipeline development, and task implementation, and to Partha Sarathi Mohanty and Riddhi Bhalerao for support with saliva sample processing. We also thank Priyanshu, Mrugsen Nagsen Gopnarayan, and Avisha for assistance with data collection.

## Author Contributions

P.M. and A.R. designed the experiment. A.R. and P.M. collected the data. P.M. performed model analysis and analyzed the behavioral data. G.C. and P.M. analyzed the salivary data. P.M. analyzed the pupil data. P.M. and A.R. wrote the manuscript. M.L.P. and A.R. provided study resources. All authors revised the manuscript.

