## supplementary file for "Trait anxiety drives premature disengagement despite intact opportunity-cost sensitivity"

Arjun Ramakrishnan

Department of Biological Sciences and Bioengineering,

Indian Institute of Technology Kanpur

Kanpur, Uttar Pradesh,208016

### Supplementary Note from Introduction

People with anxiety often disengage too quickly from tasks or opportunities, even when doing so reduces potential rewards and wellbeing. Why this occurs remains unclear. Using a naturalistic explore–exploit decision-making task, we show that anxious individuals still understand when it is beneficial to persist in a rewarding environment, yet nevertheless leave earlier than people without anxiety. Computational modeling revealed that this behavior arises because anxiety dampens the accumulation of reward information that drives decisions. Measurements of pupil size and stress-related biomarkers further showed that differences in physiological arousal shape this decision process. Together, these findings suggest that anxiety does not distort how people perceive rewards, but instead alters how reward information is translated into sustained engagement. By linking behavior, computational mechanisms, and physiological signals, this work identifies premature disengagement as a key mechanism through which anxiety can impair everyday life choices.

### Supplementary Methods

#### *Participants*

US cohort. Data for the US cohort were collected at the Wharton Behavioral Laboratory (University of Pennsylvania). All participants were in good health and reported no use of medications that could affect cognitive performance. The study was approved by the Institutional Review Board of the University of Pennsylvania, and all participants provided informed consent prior to participation. A total of 137 healthy individuals (40 males, 97 females; age range 19–45 years) were included. Each participant completed a single experimental session lasting approximately 20 minutes, during which they performed a sequential patch-foraging task comprising ~180–200 trials.

Indian cohort. Data for the Indian cohort were collected in a controlled laboratory setting at the Indian Institute of Technology Kanpur. Participants were recruited from the student population. To minimize circadian variability and ensure consistency for salivary biomarker collection, all sessions were conducted between 12:00 PM and 7:00 PM. All participants were in good health and reported no use of medications that could affect cognitive performance. The study was approved by the Institute Human Ethics Committee (IHEC) of IIT Kanpur (IITK/IEC/2020-21/II/10), and written informed consent was obtained from all participants prior to participation. A total of 55 healthy individuals (37 males, 18 females; age range 18–32 years) participated. Each participant completed a single experimental session lasting approximately one hour, including a 10-minute sequential patch-foraging task (~180–200 trials, depending on response times) along with simultaneous electroencephalography (EEG) recording, eye-tracking, and saliva sample collection.

***Exclusion criteria***

US cohort. An initial sample of 137 participants was recruited for the behavioral task. Following preprocessing, quality-control criteria were applied to ensure reliability of both self-report and behavioral data. First, 58 participants were excluded due to incomplete or invalid responses on the State–Trait Anxiety Inventory (STAI), including missing items or total scores falling below the theoretical minimum of 20. Second, to ensure adequate task engagement, participants exhibiting extreme behavioral outliers were excluded; specifically, individuals with mean handling times exceeding the 75th percentile of the sample distribution were removed, as this pattern suggested atypical or disengaged task performance. After applying these criteria, the final sample comprised 79 participants for behavioral analyses, including patch reward regression and stay duration models (which additionally incorporated trial-level exclusions), and 123 participants for hierarchical sequential sampling model (HSSM) analyses.

Indian cohort. Following preprocessing, data from eight participants were excluded based on predefined quality-control criteria, including failure to pass embedded attention checks, incomplete STAI responses, and insufficient task engagement. Engagement was operationalized as completing at least 25 trials within an environment, as valid patch-level estimates required at least one exploitation choice per patch. Participants who did not follow task instructions were also excluded. In addition, to ensure completeness of multimodal measurements, participants with missing salivary biomarker data or pupil recordings were removed from the final dataset. After applying these criteria, the remaining sample consisted of participants with complete behavioral, neural (EEG), and physiological (pupil, salivary) data, ensuring consistency and reliability across analyses.

***Foraging game***

Paradigm. To examine how trait anxiety influences sequential foraging decisions, participants performed a computer-based patch-foraging task (Fig. 1A–C), adapted from established paradigms (Addicott et al., 2017; Constantino & Daw, 2015; Hayden et al., 2011; Ramakrishnan, Pardes, et al., 2019). Participants were first presented with written instructions and completed a practice block to familiarize themselves with the task. The practice block matched the structure of the main task but used different parameters (e.g., altered travel durations and reward schedules) to ensure learning without contaminating experimental behavior. Participants were explicitly instructed that their monetary bonus was proportional to the total number of berries collected, incentivizing reward maximization.

Task structure. On each trial (Fig. 1C), participants decided whether to continue harvesting from the current patch or leave to search for a new patch. Decisions were implemented through a two-step fixation procedure designed to isolate a pre-decisional (tonic) period. Participants were required to position the cursor within a central fixation box and make an initial click, followed by a 1-s waiting period. Once the fixation cue turned dark green, a second click confirmed readiness to act. Premature movements or failure to complete the sequence reset the trial. Following successful fixation, participants could either move to the current patch to harvest or move to an exit cue (a hollow square with an upward arrow) to leave the patch. Harvesting yielded a reward displayed upon clicking the patch. Leaving initiated a transition to a new patch, with the screen scrolling downward to simulate forward movement. The duration of this transition, termed travel time, varied across task blocks and indexed environmental opportunity cost.

Reward structure. For the US cohort, the initial reward at each patch was given by $r0 = 7 + N(0, 0.25)$, and rewards decreased with each harvest according to a stochastic depletion function: $ri = ri-1 - 0.5 + N(0, 0.25)$. Rewards were bounded at zero ($ri \geq 0$), such that depleted patches yielded no further returns. The task consisted of two 4-minute blocks with different travel times: a short block (5 s) and a long block (20 s). For the Indian cohort, initial patch rewards were uniformly sampled between 85 and 100. Subsequent harvests reduced rewards by a fixed decrement ($\varepsilon = 10$) with added stochasticity: $Ri = rand(R_{i-1}^{upper} - \varepsilon, R_{i-1}^{lower} - \varepsilon)$, with rewards bounded at zero. The task consisted of two 5-minute blocks with short (3 s) and long (10 s) travel times.

Participants were not explicitly informed about travel times, requiring them to learn environmental structure through experience. Total reward (berries collected) was displayed at the end of each block. Task visuals—including the central fixation cue, transition arrow, and patch icons—were held constant across conditions. The task was implemented in PsychoPy and presented on a 27-inch LED monitor (1920 × 1080 resolution, 60 Hz refresh rate).

#### ***Self-report surveys***

At the beginning of the session, participants were provided with a study description and informed consent form, followed by collection of demographic information (age and gender). Participants then completed the Spielberger State–Trait Anxiety Inventory (STAI), including both the state (STAI-Y1) and trait (STAI-Y2) subscales; each subscale consists of 20 items rated on a 4-point Likert scale. State anxiety (STAI-Y1) was assessed twice—once prior to the task and once immediately after task completion—to capture potential changes in momentary anxiety associated with task engagement. Trait anxiety (STAI-Y2), indexing a stable disposition to experience anxiety, was assessed once at the beginning of the session.

***Pupil data acquisition, preprocessing, and epoch extraction***

Pupil diameter was recorded using a Pupil Core eye tracker (Pupil Labs GmbH, Germany) at 200 Hz. A 2D, 5-point calibration was performed at the start of each session. Eye-tracking data were streamed via Lab Streaming Layer (LSL) to ensure precise synchronization with behavioral and EEG event markers. Analyses were restricted to the left eye to maintain consistency across participants. Preprocessing was conducted using the PyPLR toolbox. Blink artifacts were identified based on rapid changes in pupil size (thresholded on the first derivative) and removed. Missing segments due to blinks were linearly interpolated. The resulting time series was smoothed using a rolling-mean filter to reduce high-frequency noise while preserving task-relevant fluctuations in pupil-linked arousal.

To dissociate sustained (tonic) and event-related (phasic) components of pupil-linked arousal, epochs were extracted relative to decision events. Tonic pupil size was defined as the mean pupil diameter within a pre-decisional window spanning −750 ms to −250 ms relative to the center-cue click; this window captures baseline arousal preceding choice commitment and reflects ongoing neuromodulatory state. Phasic pupil responses were quantified within a post-decision window from 1000 ms to 1250 ms following the decision, capturing transient pupil dilation associated with reward processing and decision execution.

***Behavioral analysis***

Behavioral analyses focused on characterizing patch-leaving decisions and reward accumulation in a sequential foraging context. Key measures included patch stay duration (time spent exploiting a patch before leaving), handling time (reaction time plus enforced waiting period), and travel time (fixed at 3 or 10 s in the Indian setup, and 5 or 20 s in the US setup, depending on environment), which operationalized environmental opportunity cost. Reward-related measures included per-trial reward, cumulative reward within a patch (indexing reward accumulation), total reward across the task, and average reward rate within each environment. These metrics quantified how efficiently participants accumulated reward over time. Trait anxiety was indexed using STAI-Y2 scores. To assess decision optimality, behavior was compared against predictions from the Marginal Value Theorem (MVT), which specifies the optimal patch-leaving policy as a function of opportunity cost. Deviations from MVT predictions—manifesting as premature patch leaving or reduced reward accumulation—were used to quantify suboptimal foraging behavior.

#### ***Deviation from optimal foraging***

To quantify decision optimality in the foraging task, we compared participants' behavior against a normative model derived from the MVT. This framework specifies the optimal number of harvests within a patch as a function of reward depletion and environmental opportunity cost (indexed by travel time and individual response dynamics). For a given patch, the cumulative reward obtained by harvesting x times was computed from the sequentially decreasing reward structure: $R_{patch}(x) = \Sigma\ (i = 1 \text{ to } x)\ r_i$, where $r_i$ denotes the reward obtained on the i-th harvest.

At the environment level, total reward accumulation depends on both the reward obtained per patch and the number of patches visited, which is constrained by the total time available. Critically, the number of patches an individual can exploit is determined by their handling time (reaction time plus enforced delay) and the travel time between patches, which together define the opportunity cost of continued exploitation. For each participant and environment (short vs long travel time), we estimated the optimal harvest number x* that maximizes the reward accumulation rate; this optimal policy reflects the trade-off between continued exploitation of a depleting patch and leaving to search for a new one. We then compared this normative benchmark to observed behavior to quantify suboptimality, computing the patch-wise reward deviation as the difference between the participant's actual cumulative reward within a patch and the model-predicted optimal reward. This measure captured deviations from optimal foraging, including premature patch leaving and reduced reward accumulation, and was used to relate individual differences in behavior to trait anxiety and neuromodulatory signals.

#### *Statistical analysis*

All statistical analyses were performed in R (version 4.4.1). We used a combination of Bayesian regression models implemented in brms, mediation analyses, and hierarchical drift diffusion / sequential sampling models to examine relationships between behavioral measures (patch leaving and reward accumulation), physiological indices (pupil dynamics, salivary markers), and individual differences in trait anxiety.

Bayesian regression models. Given the non-Gaussian structure of several behavioral and physiological measures, we employed Bayesian regression models (BRMs) in addition to complementary frequentist analyses. BRMs were particularly suited for capturing hierarchical structure and potential nonlinear relationships. Models were implemented using brms, with visualization and posterior checks performed using bayesplot. Behavioral models examined how patch leaving and reward accumulation varied as a function of environmental opportunity cost and trait anxiety:

- *Patch stay duration ~ travel time × trait anxiety*
- *Patch-wise cumulative reward ~ travel time × trait anxiety*

Pupil models were specified at the trial level with random intercepts for participants:

- *Phasic pupil response ~ reward + (1 | subject)*
- *Phasic pupil response ~ average reward + cumulative reward + travel time + (1 | subject)*
- *Phasic pupil response ~ average reward + cumulative reward + time elapsed + (1 | subject)*
- *Tonic pupil size ~ travel time + average reward + time elapsed + (1 | subject)*

To examine individual differences, we additionally modeled direct and interaction effects of trait anxiety and neuromodulatory signals:

- *Phasic pupil response ~ trait anxiety*
- *Tonic pupil size ~ trait anxiety*
- *Phasic pupil response ~ trait anxiety × baseline cortisol*
- *Phasic pupil response ~ trait anxiety × baseline salivary α-amylase (sAA)*

Mediation model. To assess whether pupil-linked arousal statistically accounted for the relationship between trait anxiety and behavioral suboptimality, we conducted mediation analyses testing whether pupil measures (e.g., phasic pupil responses) mediated the association between trait anxiety and reward accumulation (cumulative reward or average reward rate). Mediation models were implemented using the mediation and lavaan packages in R. Indirect effects were estimated using a quasi-Bayesian Monte Carlo approach with 500 simulations to obtain confidence intervals. A representative model was specified as: mediator model, phasic pupil response ~ trait anxiety; outcome model, cumulative reward ~ phasic pupil response + trait anxiety. These analyses were interpreted conservatively as statistical mediation, providing evidence for potential intermediate mechanisms linking trait anxiety, pupil-linked arousal, and suboptimal foraging behavior without implying causal directionality.

Hierarchical Sequential Sampling Modeling (HSSM). To characterize the latent decision processes underlying trial-level foraging choices (stay vs leave), we employed Hierarchical Sequential Sampling Models (HSSM), a Bayesian framework for modeling evidence accumulation in two-alternative decisions (Fengler et al.). These models assume that decisions arise from a stochastic process in which evidence favoring competing options is accumulated over time until a decision boundary is reached; in the foraging task, evidence accumulates toward one of two boundaries corresponding to continued exploitation (stay) or patch leaving (leave). Within this framework, the drift rate (v) reflects the rate at which reward-related evidence is accumulated and indexes sensitivity to reward information; the boundary separation (a) represents the amount of evidence required to commit to a decision and reflects decision caution; and the starting point (z) captures any initial bias toward staying or leaving. Reaction times and choices were jointly modeled to estimate these parameters. The hierarchical structure allows parameters to be estimated at both group and participant levels, improving statistical efficiency while accounting for individual variability. Trial-wise predictors were incorporated to examine how reward signals, environmental opportunity cost (travel time), pupil-linked arousal, and trait anxiety modulate latent decision dynamics. Modeling was implemented using HSSM (v0.2.11), with supporting libraries including Bambi (v0.15.0), NumPyro, PyMC (≥5.25), and ArviZ (≥0.22.0) for Bayesian inference and model diagnostics.

Model specifications. We evaluated a series of models of increasing complexity:

- *Model 1 (behavioral): v ~ reward + travel time + trait anxiety;  a ~ reward + travel time + trait anxiety.*
- *Model 2 (pupil-constrained): v ~ reward + travel time + phasic pupil + trait anxiety;  a ~ reward + travel time + phasic pupil + trait anxiety.*
- *Model 3 (neuromodulatory): v ~ reward + travel time + phasic pupil + trait anxiety + baseline sAA + baseline cortisol;  a ~ reward + travel time + phasic pupil + trait anxiety + baseline sAA + baseline cortisol.*

These models allowed us to test how reward-related evidence accumulation and decision thresholds are jointly shaped by environmental structure, individual differences in anxiety, and physiological indices of neuromodulatory state.

#### *Salivary cortisol and α-amylase*

Saliva samples were collected at two time points: baseline (T1, pre-task) and post-task (T2, following the foraging task). Samples were obtained using Salivette Cortisol Kits (Sarstedt), centrifuged at 3000 × g for 10 minutes, and the supernatant was aliquoted and stored at −80 °C until analysis. Cortisol concentrations were quantified using a commercially available ELISA kit (ELK8029; ELK Biotechnology, China), with a sensitivity of 0.93 ng/mL and a detection range of 3.13–200 ng/mL. Salivary α-amylase (sAA) levels were measured using an ELISA kit (E-EL-H0320; Elabscience), with a sensitivity of 0.94 ng/mL and a detection range of 1.56–100 ng/mL. All samples were assayed in duplicate, and dilution factors were adjusted as necessary to ensure that measurements fell within the linear range of the standard curve, minimizing assay variability and ensuring accuracy. Baseline (T1) cortisol and sAA levels were used as indices of individual differences in neuromodulatory tone, reflecting hypothalamic–pituitary–adrenal (HPA) axis and noradrenergic system activity, respectively. These measures were incorporated into behavioral, pupil, and HSSM analyses to examine how physiological state constrains reward-guided decision-making and patch-leaving behavior.

### Supplementary Results

#### Supplementary Figures:

Figure S1: Effect of Trait Anxiety and Travel Time on Patch Stay Duration Across Demographics.

Figure S2:  Effect of Trait Anxiety and Travel Time on Patch Reward Across Demographic Samples.

Figure S3:Variation of tonic and phasic pupil dilation with increase in trait anxiety.

Figure S4:Tonic pupil dilation mediates the effect of trait anxiety on cumulative reward.

Figure S5: Distribution of trait anxiety scores and group classification.

Figure S6:Effects of trait anxiety and SAA on phasic pupil dilation.

Figure S7:Effects of trait anxiety and cortisol on phasic pupil dilation.

### Supplementary Tables:

Table S1: Patch stay duration variation due to different variables in Indian population

Table S2: Patch stay duration variation due to different variables in US population

Table S3: Variation of total amount of reward collected in a patch due to different variables in Indian population

Table S4: Variation of total amount of reward collected in a patch due to different variables in US population

Table S5: Effect of trait anxiety,Baseline SAA,Baseline Cortisol on phasic pupil dilation (marker of phasic LC-NE activity).

Table S6: Mediation effect of phasic pupil dilation on trait anxiety driven variation of foraging decision parameters (cumulative reward per patch).

Table S7:Mediation effect of tonic pupil dilation on trait anxiety driven variation of foraging decision parameters (cumulative reward per patch).

Table S8: Subject-level analysis of mediation effect of phasic pupil dilation on trait-anxiety-driven variation of foraging decision parameters (cumulative reward per patch).

Table S9: Effect of trait anxiety, handling time, travel time and their interaction on stay duration in a patch, analysed using bayesian regression modeling.

Table S10: Effect of trait anxiety on decision mechanism in Indian population.

Table S11: Effect of foraging parameters on decision mechanism in Indian population.

Table S12: Effect of foraging parameters and trait anxiety on decision mechanism in Indian population.

Table S13: Effect of foraging parameters ,trait anxiety,and marker of LC-NE phasic activity on decision mechanism in Indian population.

Table S14: Effect of foraging parameters ,trait anxiety, stress neuromodulators, and marker of LC-NE phasic activity on decision mechanism in Indian population.

Table S15: Effect of foraging parameters, trait anxiety, and marker of tonic LC-NE activity on decision mechanism in Indian population.

Table S16: Model comparison table for HSSM in Indian population

Table S17: Effect of trait anxiety on decision mechanism in US population.

Table S18: Effect of foraging parameters on decision mechanism in US population.

Table S19: Effect of foraging parameters and trait anxiety on decision mechanism in US population.

Table S20:Model comparison table for HSSM in US population

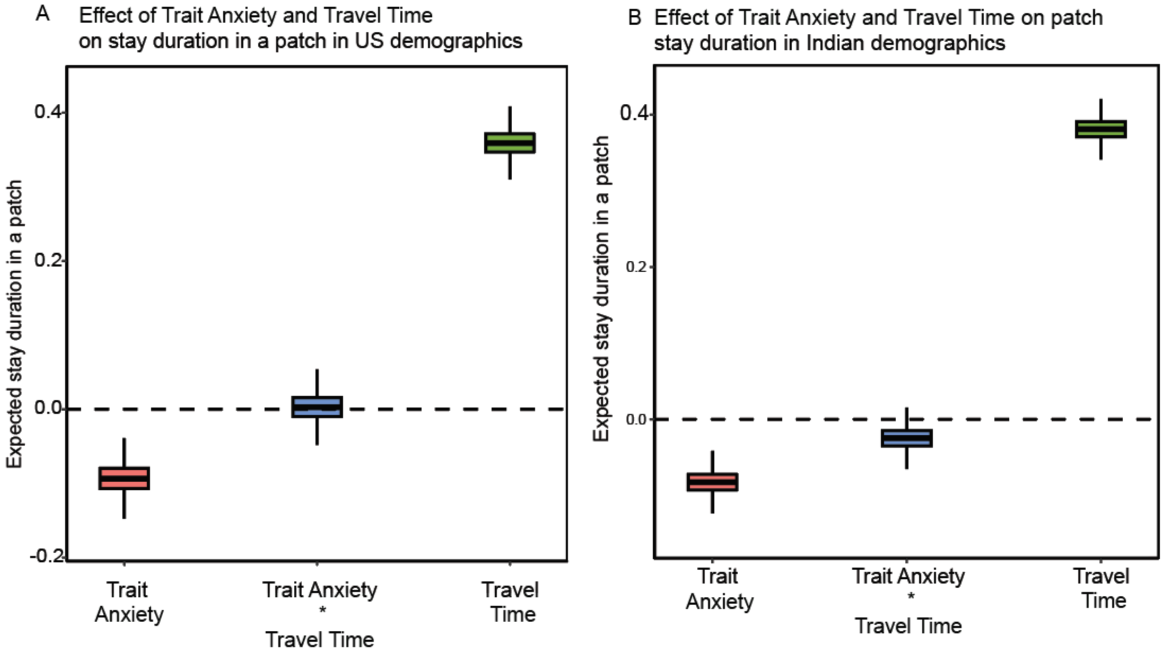


**Supplementary Figure S1. Effect of Trait Anxiety and Travel Time on Patch Stay Duration Across Demographics.**

(A) Effect of trait anxiety, travel time, and their interaction on expected stay duration within a patch in the US sample.
(B) Corresponding effects in the Indian sample.

The figure illustrates posterior distributions of regression coefficients derived from Bayesian beta regression models (logit link) predicting stay duration in a patch as a function of trait anxiety, travel time, and their interaction. Colored boxplots represent the distribution of posterior estimates for each predictor (trait anxiety, interaction term, and travel time). The central line within each box denotes the median posterior estimate, the box represents the interquartile range, and whiskers indicate the spread of the posterior distribution. The horizontal dashed line at zero denotes no effect. Coefficients above zero indicate an increase in expected stay duration associated with the predictor, whereas coefficients below zero indicate a reduction in stay duration.

Across both demographic samples, travel time shows a strong positive association with patch stay duration, consistent with adaptive foraging predictions. Trait anxiety is associated with reduced stay duration (negative coefficients), whereas the interaction between trait anxiety and travel time remains comparatively small, suggesting limited modulation of travel-time sensitivity by anxiety levels. Differences in effect magnitude between panels reflect demographic variation in behavioral sensitivity to travel costs.

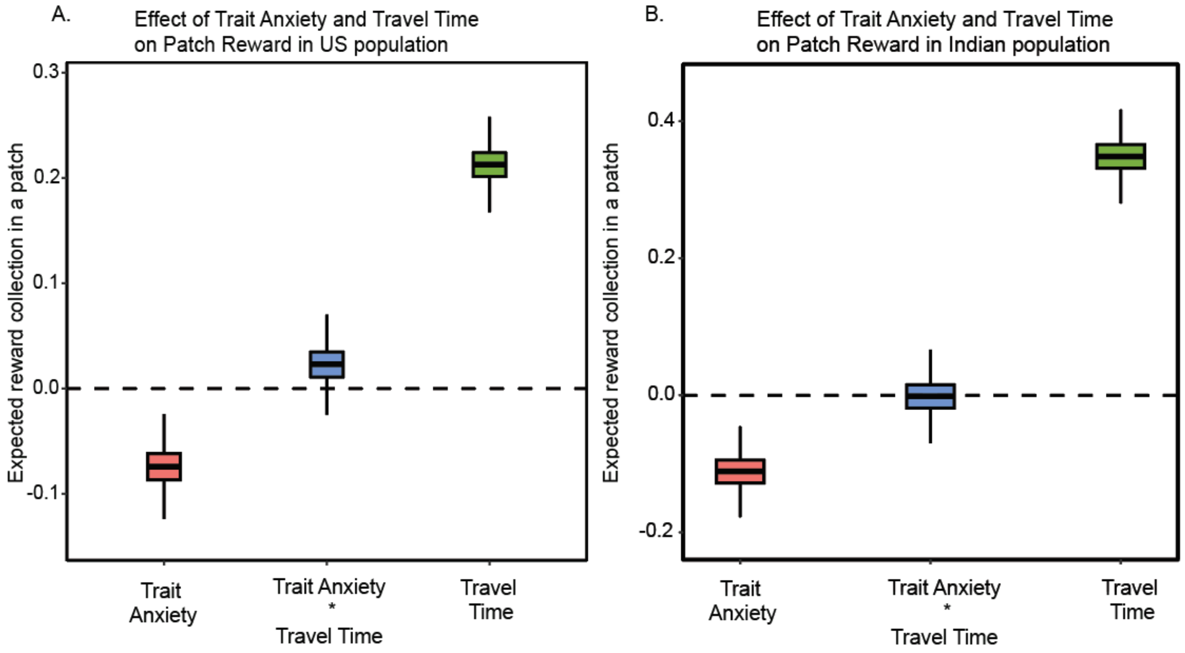


**Supplementary Figure S2. Effect of Trait Anxiety and Travel Time on Patch Reward Across Demographic Samples.**

(A) Effect of trait anxiety, travel time, and their interaction on expected patch reward in the US population.
(B) Corresponding effects in the Indian population.

The figure illustrates posterior distributions of regression coefficients derived from Bayesian beta regression models (logit link) predicting patch reward as a function of trait anxiety, travel time, and their interaction. Colored boxplots represent the distribution of posterior estimates for each predictor (trait anxiety, interaction term, and travel time). The central horizontal line within each box denotes the median posterior estimate, the box represents the interquartile range, and whiskers indicate the dispersion of the posterior distribution. The horizontal dashed line at zero denotes no effect. Coefficients above zero indicate an increase in expected reward collection within a patch associated with the predictor, whereas coefficients below zero indicate reduced reward collection.

Across both demographic samples, travel time exhibits a strong positive association with patch reward, consistent with adaptive foraging theory, whereby higher travel costs promote longer exploitation and increased reward accumulation. Trait anxiety is associated with reduced reward collection (negative coefficients), suggesting that higher anxiety relates to shorter or less efficient patch exploitation. The interaction term remains comparatively small in magnitude, indicating limited modulation of travel-cost sensitivity by anxiety levels. Differences in effect magnitude between panels reflect demographic variation in behavioral sensitivity to environmental travel costs.

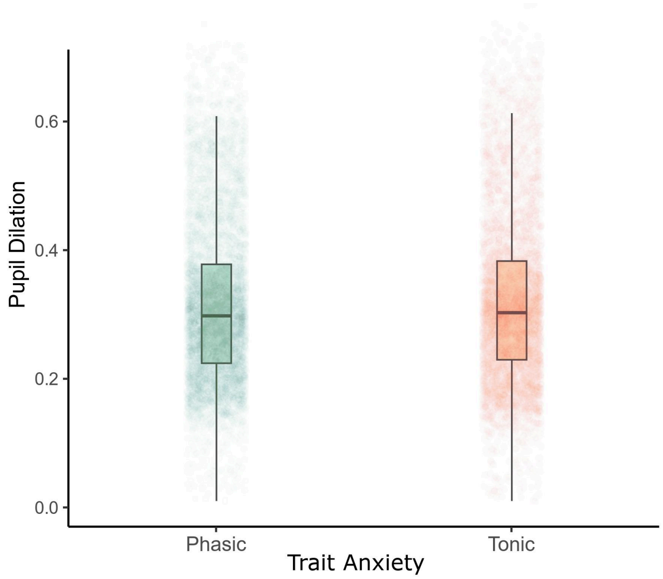


**Supplementary Figure S3. Variation of tonic and phasic pupil dilation with increase in trait anxiety.**

Posterior estimates from a Bayesian Gamma regression (log link) indicate a strong positive association between standardized trait anxiety and phasic ($\beta = 0.72$, 95% CrI [0.65, 0.79]) and tonic($\beta = 0.69$, 95% CrI [0.61, 0.76]) pupil responses. Higher trait anxiety reliably predicts increased phasic and tonic pupil dilation, consistent with heightened transient arousal responses.

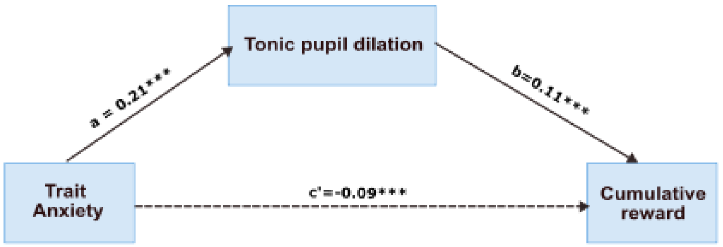


**Supplementary Figure S4 Tonic pupil dilation mediates the effect of trait anxiety on reward.**

Mediation model illustrating the indirect effect of trait anxiety on cumulative reward via tonic pupil dilation. Solid arrows indicate significant paths with standardized coefficients shown (*a* and *b*), while the dashed arrow denotes the direct effect of trait anxiety on cumulative reward (*c′*).

Path coefficients represent standardized estimates. Solid arrows denote significant effects; dashed arrows denote direct effects. *** denotes significance level $p < 0.001$.

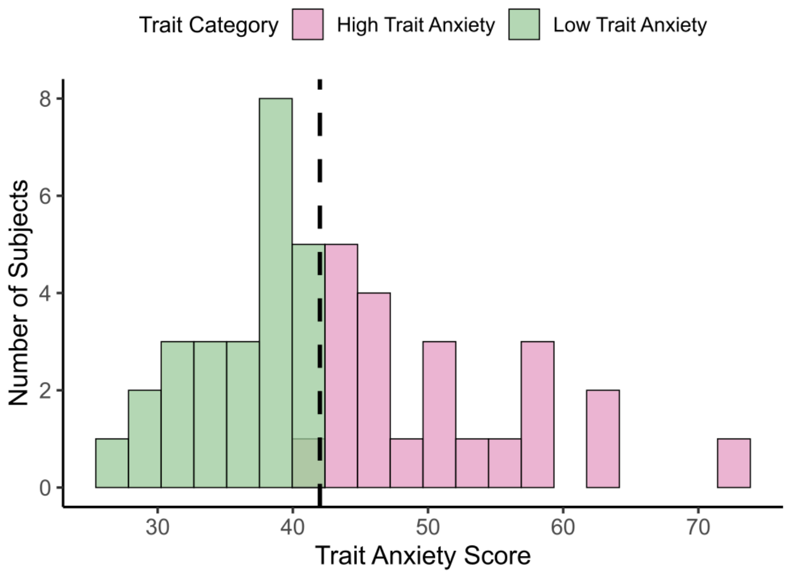


**Supplementary Figure S5. Distribution of trait anxiety scores and group classification.**

Histogram showing the distribution of trait anxiety scores across participants. Individuals were categorized into low trait anxiety (green) and high trait anxiety (pink) groups based on a median split (dashed vertical line). The y-axis denotes the number of subjects per bin.

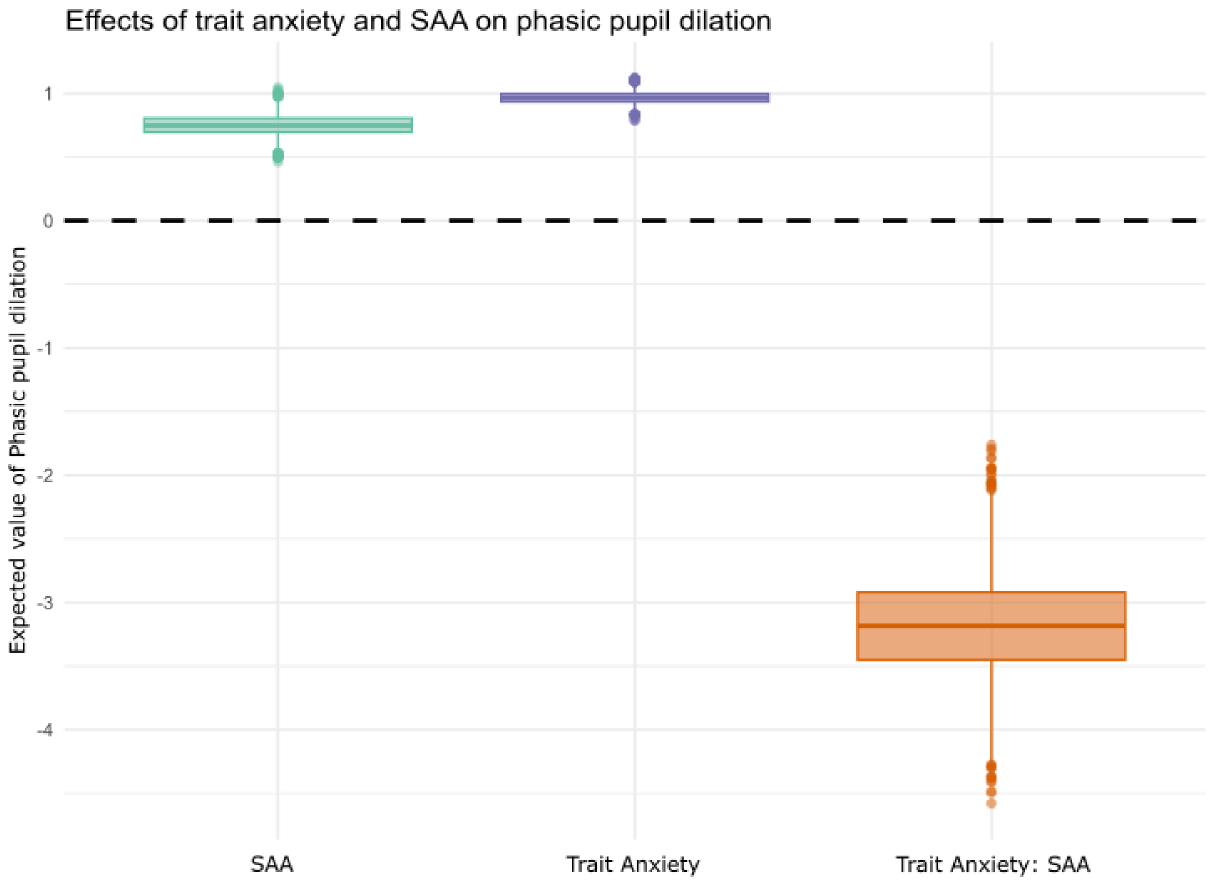


**Supplementary Figure S6.Effects of trait anxiety and SAA on phasic pupil dilation.**

Boxplots show posterior expected values of phasic pupil dilation for the main effects of salivary alpha-amylase (SAA), trait anxiety, and their interaction. Boxes indicate the interquartile range, center lines denote the median, and points represent individual posterior draws. The dashed horizontal line denotes the zero reference.

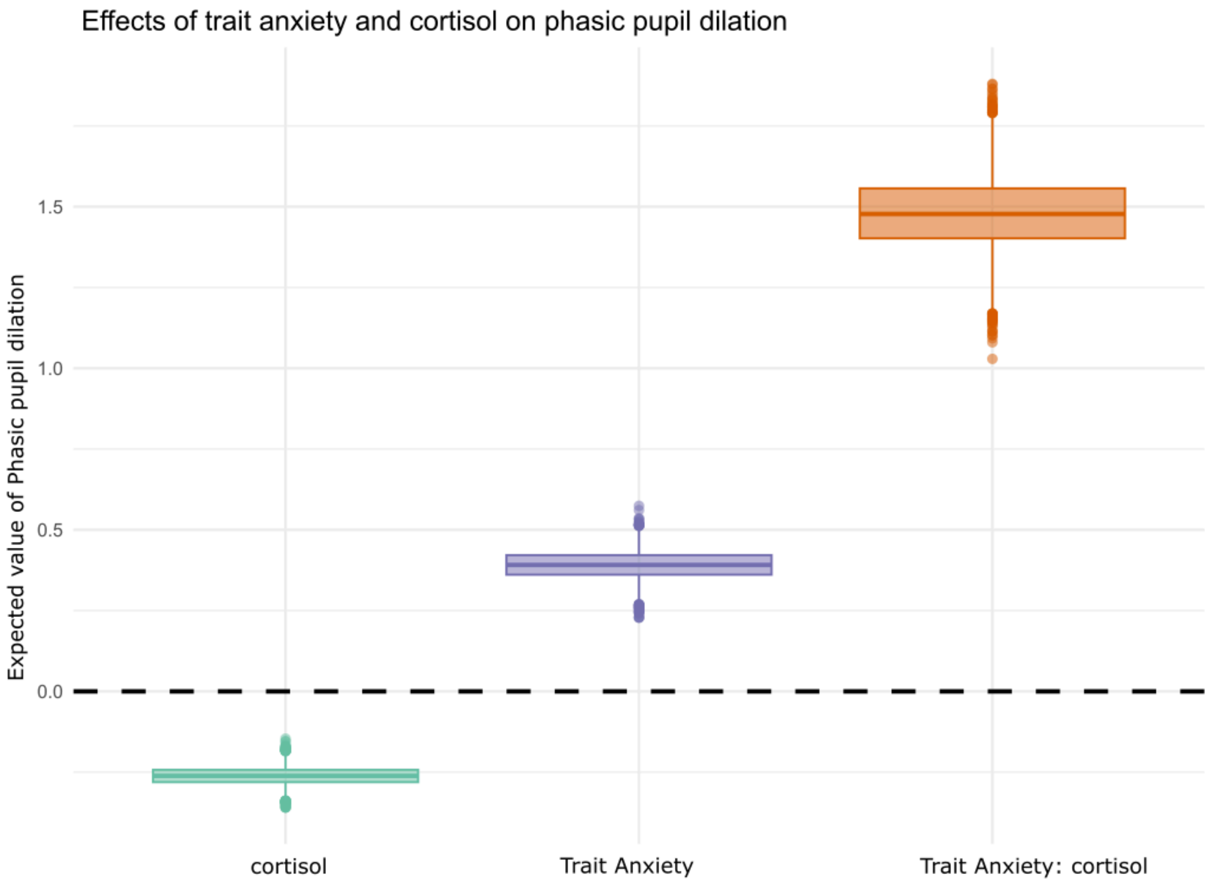


**Supplementary Figure S7. Effects of trait anxiety and cortisol on phasic pupil dilation.**

Boxplots show posterior expected values of phasic pupil dilation for the main effects of **cortisol**, **trait anxiety**, and their interaction. Boxes indicate the interquartile range, center lines denote the median, and points represent individual posterior draws. The dashed horizontal line denotes the zero reference.

**Supplementary tables:**

**Table S1. Patch stay duration variation due to different variables in Indian population**

| 1. Equation : Patch stay duration ~ Travel time*Trait anxiety (Bayesian Regression modelling for Indian demography) | | | | | | | |
|---|---|---|---|---|---|---|---|
| **Parameter** | **Estimate** | **Est. Error** | **95% CI (Lower)** | **95% CI (Upper)** | **Rhat** | **Bulk_ESS** | **Tail_ESS** |
| **Intercept** | -0.50 | 0.02 | -0.53 | -0.47 | 1.00 | 12371 | 8180 |
| **Trait_anxiety** | -0.08 | 0.02 | -0.11 | -0.05 | 1.00 | 12668 | 8623 |
| **Travel_time** | 0.38 | 0.01 | 0.35 | 0.41 | 1.00 | 12057 | 8041 |
| **Trait_anxiety × Travel_time** | -0.02 | 0.01 | -0.05 | 0.00 | 1.00 | 12333 | 8246 |
| **Phi** | 14.16 | 0.54 | 13.11 | 15.22 | 1.00 | 12366 | 7838 |

**Table S2. Patch stay duration variation due to different variables in US population**

| 2. Equation : Patch stay duration ~ Travel time*Trait anxiety (Bayesian Regression modelling for US demography) | | | | | | | |
|---|---|---|---|---|---|---|---|
| **Parameter** | **Estimate** | **Est. Error** | **95% CI (Lower)** | **95% CI (Upper)** | **Rhat** | **Bulk_ESS** | **Tail_ESS** |
| **Intercept** | -0.64 | 0.02 | -0.68 | -0.61 | 1.00 | 14224 | 7977 |
| **Trait_anxiety** | -0.09 | 0.02 | -0.13 | -0.06 | 1.00 | 14378 | 7928 |
| **Travel_time** | 0.36 | 0.02 | 0.32 | 0.40 | 1.00 | 13216 | 7177 |
| **Trait_anxiety × Travel_time** | 0.00 | 0.02 | -0.04 | 0.04 | 1.00 | 13978 | 7127 |
| **Phi** | 3.79 | 0.10 | 3.60 | 3.99 | 1.00 | 13978 | 7127 |

**Table S3. Variation of total amount of reward collected in a patch due to different variables in Indian population**

| 3. Equation : Total reward gain in a patch ~ Travel time*Trait anxiety (Bayesian Regression modelling for Indian demography) | | | | | | | |
|---|---|---|---|---|---|---|---|
| **Parameter** | **Estimate** | **Est. Error** | **95% CI (Lower)** | **95% CI (Upper)** | **Rhat** | **Bulk_ESS** | **Tail_ESS** |
| **Intercept** | 0.56 | 0.03 | 0.51 | 0.61 | 1.00 | 13030 | 7870 |
| **Travel_time** | 0.35 | 0.03 | 0.30 | 0.40 | 1.00 | 14235 | 7210 |
| **Trait_anxiety** | -0.11 | 0.02 | -0.16 | -0.06 | 1.00 | 13690 | 7275 |
| **Travel_time × Trait_anxiety** | -0.00 | 0.03 | -0.05 | 0.05 | 1.00 | 14529 | 7352 |
| **Phi** | 4.30 | 0.16 | 3.99 | 4.61 | 1.00 | 13794 | 7275 |

**Table S4.Variation of total amount of reward collected in a patch due to different variables in US population**

| 4. Equation : Total reward gain in a patch ~ Travel time*Trait anxiety (Bayesian Regression modelling for US demography) | | | | | | | |
|---|---|---|---|---|---|---|---|
| **Parameter** | **Estimate** | **Est. Error** | **95% CI (Lower)** | **95% CI (Upper)** | **Rhat** | **Bulk_ESS** | **Tail_ESS** |
| **Intercept** | -1.31 | 0.02 | -1.35 | -1.27 | 1.00 | 10227 | 7665 |
| **Trait_anxiety** | -0.07 | 0.02 | -0.11 | -0.04 | 1.00 | 11936 | 7952 |
| **Travel_time** | 0.21 | 0.02 | 0.18 | 0.25 | 1.00 | 11345 | 7605 |
| **Trait_anxiety × Travel_time** | 0.02 | 0.02 | -0.01 | 0.06 | 1.00 | 12481 | 7299 |
| **Phi** | 3.79 | 0.10 | 3.60 | 3.99 | 1.00 | 11869 | 7673 |

**Supplementary Table.5 - Effect of trait anxiety, Baseline SAA, Baseline Cortisol on phasic pupil dilation (marker of phasic LC-NE activity).**

**Supplementary table 5.A**

**Equation : Pupildata_phasic ~ Trait_anxiety (Bayesian regression modeling)**

**family = Gamma(link = "log"), chains = 4, iter = 5000, warmup = 1000**

| Predictor | Estimate | SE | 95% CI (Lower) | 95% CI (Upper) | R̂ | Bulk ESS | Tail ESS |
|---|---|---|---|---|---|---|---|
| **Intercept** | −1.33 | 0.01 | −1.35 | −1.31 | 1.00 | 13,878 | 11,535 |
| **Trait anxiety (z)** | 0.72 | 0.04 | 0.65 | 0.79 | 1.00 | 13,112 | 11,438 |

**Supplementary table 5.B**

**Equation : Pupildata_phasic ~ Baseline SAA*Trait_anxiety (Bayesian Regression modelling)**

**family = Gamma(link = "log"), chains = 4, iter = 5000, warmup = 1000**

| Predictor | Estimate | SE | 95% CI (Lower) | 95% CI (Upper) | R̂ | Bulk ESS | Tail ESS |
|---|---|---|---|---|---|---|---|
| **Intercept** | −1.40 | 0.01 | −1.42 | −1.37 | 1.00 | 10,820 | 10,513 |
| **Baseline SAA (z)** | 0.75 | 0.08 | 0.60 | 0.90 | 1.00 | 7,395 | 8,444 |
| **Trait anxiety (z)** | 0.96 | 0.04 | 0.88 | 1.05 | 1.00 | 9,550 | 9,396 |
| **Baseline SAA × Trait anxiety (z)** | −3.19 | 0.38 | −3.92 | −2.46 | 1.00 | 7,393 | 8,468 |

**Supplementary table 5.C**

**Equation : Pupildata_phasic ~ Baseline cortisol*Trait_anxiety (Bayesian Regression modelling)**

**family = Gamma(link = "log"), chains = 4, iter = 5000, warmup = 1000**

| Predictor | Estimate | SE | 95% CI (Lower) | 95% CI (Upper) | $\hat{R}$ | Bulk ESS | Tail ESS |
|---|---|---|---|---|---|---|---|
| **Intercept** | −1.27 | 0.01 | −1.29 | −1.24 | 1.00 | 10,285 | 11,202 |
| **Baseline cortisol (z)** | −0.26 | 0.03 | −0.32 | −0.21 | 1.00 | 7,521 | 9,376 |
| **Trait anxiety (z)** | 0.39 | 0.04 | 0.31 | 0.48 | 1.00 | 9,632 | 10,360 |
| **Baseline cortisol × Trait anxiety (z)** | 1.48 | 0.11 | 1.26 | 1.70 | 1.00 | 7,304 | 8,996 |

**Supplementary Table S6 - Mediation effect of phasic pupil dilation on trait anxiety driven variation of foraging decision parameters (cumulative reward per patch).**

**Supplementary table.6A**
**Mediator model:**
*Phasic pupil dilation* ~ Trait anxiety

| Outcome (Mediator) | Predictor | Estimate | SE | t | p |
|---|---|---|---|---|---|
| **Phasic pupil dilation** | **Intercept** | 0.000 | 0.012 | 0.00 | 1.000 |
| **Phasic pupil dilation** | **Trait anxiety (*a* path)** | 0.226 | 0.012 | 19.54 | < 0.001 |

**Supplementary table.6B**
**Outcome model:**
*Cumulative reward* ~ Phasic pupil dilation + Trait anxiety

| Outcome | Predictor | Estimate | SE | t | p |
|---|---|---|---|---|---|
| **Cumulative reward** | **Intercept** | 0.000 | 0.012 | 0.00 | 1.000 |
| **Cumulative reward** | **Phasic pupil dilation (*b* path)** | 0.104 | 0.012 | 8.59 | < 0.001 |
| **Cumulative reward** | **Trait anxiety (*c′* path)** | −0.092 | 0.012 | −7.58 | < 0.001 |

**Supplementary table.6C**

| Causal mediation effects (quasi-Bayesian, 500 simulations) | | | | |
|---|---|---|---|---|
| **Effect** | **Estimate** | **95% CI Lower** | **95% CI Upper** | **p** |
| **ACME (Indirect effect)** | 0.024 | 0.018 | 0.029 | < 0.001 |
| **ADE (Direct effect)** | −0.091 | −0.114 | −0.067 | < 0.001 |
| **Total effect** | −0.068 | −0.090 | −0.044 | < 0.001 |
| **Proportion mediated** | −0.347 | −0.545 | −0.236 | < 0.001 |

**All variables were standardized prior to analysis.**
**ACME = average causal mediation effect; ADE = average direct effect.**
**Confidence intervals were estimated using quasi-Bayesian bootstrapping (500 simulations).**
**Sample size: N = 7,069**

**Supplementary table S7 -Mediation effect of tonic pupil dilation on trait anxiety driven variation of foraging decision parameters (cumulative reward per patch).**

| **Supplementary table.7A**<br>Regression models (trial-wise)<br>**Mediator model:**<br>*Tonic pupil dilation* ~ Trait anxiety | | | | | |
|---|---|---|---|---|---|
| **Outcome (Mediator)** | **Predictor** | **Estimate** | **SE** | **t** | **p** |
| **Tonic pupil dilation** | **Intercept** | 0.000 | 0.012 | 0.00 | 1.000 |
| **Tonic pupil dilation** | **Trait anxiety (*a* path)** | 0.213 | 0.012 | 18.29 | < 0.001 |

| **Supplementary table.7B**<br>**Outcome model:**<br>*Cumulative reward* ~ Tonic pupil dilation + Trait anxiety | | | | | |
|---|---|---|---|---|---|
| **Outcome** | **Predictor** | **Estimate** | **SE** | **t** | **p** |
| **Cumulative reward** | **Intercept** | 0.000 | 0.012 | 0.00 | 1.000 |
| **Cumulative reward** | **Tonic pupil dilation (*b* path)** | 0.107 | 0.012 | 8.83 | < 0.001 |

| | | | | | |
|---|---|---|---|---|---|
| **Cumulative reward** | **Trait anxiety (*c′* path)** | −0.091 | 0.012 | −7.54 | < 0.001 |

| **Supplementary table.7C**<br>**Causal mediation effects (trial-wise, quasi-Bayesian, 500 simulations)** | | | | |
|---|---|---|---|---|
| **Effect** | **Estimate** | **95% CI Lower** | **95% CI Upper** | **p** |
| **ACME (Indirect effect)** | 0.023 | 0.018 | 0.028 | < 0.001 |
| **ADE (Direct effect)** | −0.092 | −0.115 | −0.068 | < 0.001 |
| **Total effect** | −0.069 | −0.091 | −0.046 | < 0.001 |
| **Proportion mediated** | −0.334 | −0.532 | −0.230 | < 0.001 |

**Supplementary table.8 - Subject-level analysis of mediation effect of phasic pupil dilation on trait-anxiety-driven variation of foraging decision parameters (cumulative reward per patch).**

| **Supplementary table.8A**<br><br>**Mediator model:**<br>Phasic pupil dilation ~ Trait anxiety | | | | | |
|---|---|---|---|---|---|
| **Outcome (Mediator)** | **Predictor** | **Estimate** | **S.E.** | **t-value** | **p-value** |
| **Phasic pupil dilation** | **Intercept** | 0.000 | 0.146 | 0.00 | 1.000 |
| **Phasic pupil dilation** | **Trait anxiety (a path)** | 0.208 | 0.148 | 1.41 | 0.166 |

| **Supplementary table.8B**<br>**Outcome model:**<br>*Cumulative reward* ~ Subject-wise phasic pupil dilation + Trait anxiety | | | | | |
|---|---|---|---|---|---|
| **Outcome** | **Predictor** | **Estimate** | **SE** | **t** | **p** |

| | | | | | |
|---|---|---|---|---|---|
| **Cumulative reward** | **Intercept** | 0.000 | 0.141 | 0.00 | 1.000 |
| **Cumulative reward** | **Phasic pupil dilation (SW) (*b* path)** | 0.314 | 0.146 | 2.15 | 0.037 |
| **Cumulative reward** | **Trait anxiety (*c′* path)** | −0.239 | 0.146 | −1.64 | 0.109 |

**Supplementary table.8C**

Mediation effects (subject-wise, quasi-Bayesian, 500 simulations)

| Effect | Estimate | 95% CI Lower | 95% CI Upper | p-value |
|---|---|---|---|---|
| **ACME (Indirect effect)** | 0.068 | −0.025 | 0.208 | 0.188 |
| **ADE (Direct effect)** | −0.240 | −0.493 | 0.040 | 0.088 |
| **Total effect** | −0.172 | −0.450 | 0.110 | 0.248 |
| **Proportion mediated** | −0.217 | −6.473 | 4.526 | 0.404 |

**Supplementary table.9 - Effect of trait anxiety, handling time,travel time and their interaction on stay duration in a patch, analysed using bayesian regression modeling.**

Handling time was defined as the total trial duration, comprising reaction time together with the experimentally fixed intervals (the 1-s central fixation and the inter-trial waiting period).

**9.a. Stay duration in a patch ~ Handling Time * Trait Anxiety * Travel time (Indian Population)**

| Predictor | Estimate | Est. Error | 95% CI (Lower) | 95% CI (Upper) | Rhat | Bulk ESS | Tail ESS |
|---|---|---|---|---|---|---|---|
| **Intercept** | -0.50 | 0.02 | -0.53 | -0.47 | 1.00 | 16213 | 7361 |
| **Handling Time** | -0.07 | 0.02 | -0.10 | -0.03 | 1.00 | 14075 | 8143 |
| **Trait Anxiety** | -0.07 | 0.02 | -0.10 | -0.03 | 1.00 | 13641 | 8521 |

| | | | | | | | |
|---|---|---|---|---|---|---|---|
| **Travel time** | 0.41 | 0.02 | 0.38 | 0.44 | 1.00 | 16073 | 8126 |
| **Handling Time * Trait Anxiety** | -0.06 | 0.02 | -0.10 | -0.02 | 1.00 | 12584 | 8563 |
| **Handling Time * Travel time** | -0.00 | 0.02 | -0.03 | 0.03 | 1.00 | 15070 | 8139 |
| **Trait Anxiety * Travel time** | 0.01 | 0.02 | -0.02 | 0.04 | 1.00 | 15003 | 8520 |
| **Handling Time * Trait Anxiety * Travel time** | -0.01 | 0.02 | -0.05 | 0.03 | 1.00 | 11843 | 7202 |

| **9.b. Stay duration in a patch ~ Handling Time * Trait Anxiety * Travel time (US Population)** | | | | | | | |
|---|---|---|---|---|---|---|---|
| **Predictor** | **Estimate** | **SE** | **95% CI (Lower)** | **95% CI (Upper)** | **Rhat** | **Bulk ESS** | **Tail ESS** |
| Intercept | -0.60 | 0.02 | -0.64 | -0.56 | 1.00 | 13111 | 8587 |
| Trait Anxiety | -0.09 | 0.02 | -0.13 | -0.05 | 1.00 | 13519 | 7972 |
| Handling Time | 0.41 | 0.03 | 0.35 | 0.46 | 1.00 | 11511 | 8340 |
| Travel Time | 0.38 | 0.02 | 0.34 | 0.41 | 1.00 | 11941 | 8253 |
| Trait Anxiety × Handling Time | -0.02 | 0.03 | -0.09 | 0.04 | 1.00 | 12894 | 7811 |
| Trait Anxiety × Travel Time | 0.03 | 0.02 | -0.01 | 0.06 | 1.00 | 12523 | 7414 |
| Handling Time × Travel Time | 0.06 | 0.03 | 0.00 | 0.12 | 1.00 | 10935 | 8033 |
| Trait Anxiety × Handling Time × Travel Time | 0.18 | 0.03 | 0.11 | 0.25 | 1.00 | 12623 | 8062 |

**Supplementary table.10 - Effect of trait anxiety on decision mechanism in Indian population.10. v ~ trait anxiety,**
**a ~ trait anxiety**

| Parameter | Predictor | Mean β | SD | 95% HDI (3–97%) | R̂ |
|---|---|---|---|---|---|
| **v (drift rate)** | Trait anxiety (z) | −0.009 | 0.010 | [−0.028, 0.010] | 1.000 |
| | Intercept | 1.763 | 0.014 | [1.738, 1.790] | 1.000 |
| **a (boundary separation)** | Trait anxiety (z) | −0.020 | 0.002 | [−0.023, −0.017] | 1.000 |
| | Intercept | 0.649 | 0.002 | [0.646, 0.653] | 1.000 |
| **t (non-decision time)** | — | 0.187 | 0.001 | [0.185, 0.188] | 1.000 |
| **z (starting point)** | — | 0.449 | 0.002 | [0.445, 0.454] | 1.000 |

**Supplementary table.11- Effect of foraging parameters on decision mechanism in Indian population**
**"v ~ Reward + Travel_time"**
**"a ~ Reward + Travel_time"**

| Parameter | Predictor | Mean β | SD | 95% HDI (3–97%) | R̂ |
|---|---|---|---|---|---|
| **v (drift rate)** | Reward (z) | 1.788 | 0.038 | [1.718, 1.860] | 1.000 |
| | Travel time (c) | 0.078 | 0.010 | [0.058, 0.098] | 1.000 |
| | Intercept | 1.619 | 0.050 | [1.526, 1.713] | 1.000 |
| **a (boundary separation)** | Reward (z) | 0.093 | 0.007 | [0.080, 0.106] | 1.000 |
| | Travel time (c) | −0.004 | 0.001 | [−0.006, −0.001] | 1.000 |
| | Intercept | 0.655 | 0.012 | [0.632, 0.678] | 1.000 |
| **t (non-decision time)** | — | 0.462 | 0.002 | [0.459, 0.465] | 1.000 |

| | | | | | |
|---|---|---|---|---|---|
| **z (starting point)** | — | 0.601 | 0.008 | [0.586, 0.615] | 1.000 |

**Supplementary table.12- Effect of foraging parameters and trait anxiety on decision mechanism in Indian population.**

**"v ~ Reward + Travel_time + Trait_Anxiety"**

**"a ~ Reward + Travel_time + Trait_Anxiety"**

| Parameter | Predictor | Mean β | SD | 95% HDI (3–97%) | R̂ |
|---|---|---|---|---|---|
| **v (drift rate)** | Reward (z) | 1.790 | 0.037 | [1.720, 1.859] | 1.000 |
| | Travel time (c) | 0.077 | 0.011 | [0.057, 0.097] | 1.000 |
| | Trait anxiety (z) | −0.081 | 0.033 | [−0.140, −0.017] | 1.000 |
| | Intercept | 1.610 | 0.050 | [1.514, 1.702] | 1.000 |
| **a (boundary separation)** | Reward (z) | 0.093 | 0.007 | [0.080, 0.107] | 1.000 |
| | Travel time (c) | −0.004 | 0.001 | [−0.006, −0.001] | 1.000 |
| | Trait anxiety (z) | −0.001 | 0.004 | [−0.010, 0.006] | 1.000 |
| | Intercept | 0.656 | 0.013 | [0.633, 0.681] | 1.000 |
| **t (non-decision time)** | — | 0.462 | 0.002 | [0.459, 0.465] | 1.000 |
| **z (starting point)** | — | 0.601 | 0.008 | [0.587, 0.616] | 1.000 |

**Supplementary table.13 - Effect of foraging parameters ,trait anxiety,and marker of LC-NE phasic activity on decision mechanism in Indian population.**

**"v ~ Reward + Travel_time + pupildata_phasic + Trait_Anxiety "**

**"a ~ Reward + Travel_time + pupildata_phasic+ Trait_Anxiety"**

| Parameter | Predictor | Mean β | SD | 95% HDI (3–97%) | R̂ |
|---|---|---|---|---|---|
| **v (drift rate)** | **Reward (z)** | 1.796 | 0.038 | [1.725, 1.866] | 1.000 |
| | **Travel time (c)** | 0.076 | 0.011 | [0.056, 0.096] | 1.000 |
| | **Trait anxiety (z)** | −0.081 | 0.033 | [−0.142, −0.019] | 1.000 |
| | **Pupil phasic (z)** | −0.064 | 0.034 | [−0.129, −0.001] | 1.000 |
| | **Intercept** | 1.616 | 0.051 | [1.521, 1.711] | 1.000 |
| **a (boundary separation)** | **Reward (z)** | 0.093 | 0.007 | [0.080, 0.106] | 1.000 |
| | **Travel time (c)** | −0.004 | 0.001 | [−0.006, −0.001] | 1.000 |
| | **Trait anxiety (z)** | −0.002 | 0.004 | [−0.009, 0.006] | 1.000 |
| | **Pupil phasic (z)** | 0.007 | 0.004 | [−0.001, 0.016] | 1.000 |
| | **Intercept** | 0.656 | 0.012 | [0.633, 0.680] | 1.000 |
| **t (non-decision time)** | — | 0.462 | 0.002 | [0.459, 0.465] | 1.000 |
| **z (starting point)** | **Intercept** | 0.601 | 0.008 | [0.587, 0.616] | 1.000 |

**Supplementary table.14- Effect of foraging parameters ,trait anxiety, stress neuromodulators, and marker of LC-NE phasic activity on decision mechanism in Indian population.**

**"v ~ Reward + Travel_time + pupildata_phasic + Trait_Anxiety +Pre_SAA + Pre_cort",**
**"a ~ Reward + Travel_time + pupildata_phasic+ Trait_Anxiety + Pre_SAA + Pre_cort"**

| Parameter | Predictor | Mean β | SD | 95% HDI (3–97%) | R̂ |
|---|---|---|---|---|---|
| **v (drift rate)** | Reward (z) | 1.798 | 0.038 | [1.727, 1.870] | 1.000 |
| | Travel time (c) | 0.075 | 0.011 | [0.054, 0.094] | 1.000 |
| | Trait anxiety (z) | −0.085 | 0.034 | [−0.149, −0.021] | 1.000 |
| | Pre-SAA (z) | −0.082 | 0.034 | [−0.147, −0.021] | 1.000 |

| | | | | | |
|---|---|---|---|---|---|
| | Pre-cort (z) | −0.036 | 0.035 | [−0.102, 0.029] | 1.000 |
| | Pupil phasic (z) | −0.062 | 0.034 | [−0.126, 0.002] | 1.000 |
| | Intercept | 1.619 | 0.050 | [1.521, 1.710] | 1.000 |
| **a (boundary separation)** | Reward (z) | 0.093 | 0.007 | [0.079, 0.106] | 1.000 |
| | Travel time (c) | −0.004 | 0.001 | [−0.006, −0.001] | 1.000 |
| | Trait anxiety (z) | −0.002 | 0.004 | [−0.010, 0.006] | 1.000 |
| | Pre-SAA (z) | −0.000 | 0.004 | [−0.009, 0.008] | 1.000 |
| | Pre-cort (z) | 0.007 | 0.005 | [−0.001, 0.016] | 1.000 |
| | Pupil phasic (z) | 0.008 | 0.004 | [−0.001, 0.016] | 1.000 |
| | Intercept | 0.657 | 0.013 | [0.633, 0.681] | 1.000 |
| **t (non-decision time)** | — | 0.462 | 0.002 | [0.459, 0.465] | 1.000 |
| **z (starting point)** | — | 0.601 | 0.008 | [0.586, 0.615] | 1.000 |

**Supplementary table. 15- Effect of foraging parameters, trait anxiety, and marker of tonic LC-NE activity on decision mechanisms in the Indian population.**

**Tonic HSSM result:**

**"v ~ Reward + Travel_time + pupildata_tonic + Trait anxiety."**

**“a ~ Reward + Travel_time + pupildata_tonic + Trait anxiety“**

| Parameter | Predictor | Mean β | SD | 95% HDI (3–97%) | $\hat{R}$ |
|---|---|---|---|---|---|
| v (drift rate) | **Reward (z)** | 1.775 | 0.037 | [1.704, 1.844] | 1.000 |
| | **Travel time (c)** | 0.070 | 0.010 | [0.050, 0.089] | 1.000 |
| | **Trait anxiety (z)** | −0.077 | 0.033 | [−0.141, −0.016] | 1.000 |
| | **Pupil tonic (z)** | −0.075 | 0.034 | [−0.140, −0.014] | 1.000 |
| | **Intercept** | 1.593 | 0.050 | [1.499, 1.689] | 1.000 |

| | | | | | |
|---|---|---|---|---|---|
| **a (boundary separation)** | **Reward (z)** | 0.094 | 0.007 | [0.082, 0.108] | 1.000 |
| | **Travel time (c)** | −0.003 | 0.001 | [−0.005, −0.000] | 1.000 |
| | **Trait anxiety (z)** | −0.003 | 0.004 | [−0.011, 0.005] | 1.000 |
| | **Pupil tonic (z)** | 0.002 | 0.004 | [−0.006, 0.010] | 1.000 |
| | **Intercept** | 0.660 | 0.012 | [0.637, 0.684] | 1.000 |
| **t (non-decision time)** | — | 0.462 | 0.002 | [0.459, 0.465] | 1.000 |
| **z (starting point)** | — | 0.602 | 0.008 | [0.588, 0.617] | 1.000 |

**Supplementary table.16 Model comparison table for HSSM in Indian population**

| **Parameters in model** | **devianceLOO** | **pLOO** | **devianceWAIC** | **pWAIC** |
|---|---|---|---|---|
| Trait anxiety | −1302.62 | 5.58 | −1302.62 | 5.58 |
| Reward + Travel_time | −4020.10 | 6.64 | −4020.10 | 6.64 |
| Reward + Travel_time + Trait_Anxiety | −4022.52 | 8.39 | −4022.52 | 8.39 |
| Reward + Travel_time + pupildata_phasic + Trait_Anxiety | −4027.22 | 10.39 | −4027.22 | 10.39 |
| **Reward + Travel_time + pupildata_phasic + Trait_Anxiety + Pre_SAA + Pre_cort** | **−4029.16** | **14.30** | **−4029.18** | **14.30** |

Hierarchical Sequential sampling model (HSSM) on the **US population's foraging behavioral data**:

**Supplementary table.17 : Effect of trait anxiety on decision mechanism in US population.**

**"v ~ Trait_Anxiety,"**
**"a ~ Trait_Anxiety"**

| **Parameter** | **Predictor** | **Mean β** | **SD** | **95% HDI (3–97%)** | **R̂** |
|---|---|---|---|---|---|

| | | | | | |
|---|---|---|---|---|---|
| **v (drift rate)** | Trait anxiety (z) | -0.018 | 0.012 | [-0.041, 0.005] | 1.000 |
| | Intercept | 1.838 | 0.016 | [1.808, 1.868] | 1.000 |
| **a (boundary separation)** | Trait anxiety (z) | -0.002 | 0.002 | [-0.005, 0.002] | 1.000 |
| | Intercept | 0.639 | 0.002 | [0.635, 0.644] | 1.000 |
| **t (non-decision time)** | — | 0.243 | 0.001 | [0.241, 0.245] | 1.000 |
| **z (starting point)** | Intercept | 0.442 | 0.003 | [0.437, 0.447] | 1.000 |

**Supplementary table.18: Effect of foraging parameters on decision mechanism in US population.**

**"v ~ Reward + Travel_time",**
**"a ~ Reward + Travel_time"**

| Parameter | Predictor | Mean β | SD | 95% HDI (3–97%) | $\hat{R}$ |
|---|---|---|---|---|---|
| **v (drift rate)** | Reward (z) | 1.483 | 0.014 | [1.457, 1.508] | 1.000 |
| | Travel time (c) | 0.024 | 0.002 | [0.021, 0.027] | 1.000 |
| | Intercept | 2.293 | 0.019 | [2.256, 2.328] | 1.000 |
| **a (boundary separation)** | Reward (z) | 0.172 | 0.004 | [0.163, 0.180] | 1.000 |
| | Travel time (c) | 0.001 | 0.000 | [0.000, 0.002] | 1.000 |
| | Intercept | 0.954 | 0.009 | [0.936, 0.971] | 1.000 |
| **t (non-decision time)** | — | 0.216 | 0.001 | [0.213, 0.218] | 1.000 |
| **z (starting point)** | Intercept | 0.526 | 0.004 | [0.519, 0.534] | 1.000 |

**Supplementary table.19: Effect of foraging parameters and trait anxiety on decision mechanism in US population.**

**"v ~ Reward + Travel_time + Trait_Anxiety ",**
**"a ~ Reward + Travel_time + Trait_Anxiety"**

| Parameter | Predictor | Mean β | SD | 95% HDI (3–97%) | R̂ |
|---|---|---|---|---|---|
| v (drift rate) | Reward (z) | 1.484 | 0.014 | [1.458, 1.509] | 1.000 |
| | Travel time (c) | 0.024 | 0.002 | [0.021, 0.027] | 1.000 |
| | Trait anxiety (z) | −0.033 | 0.012 | [−0.054, −0.010] | 1.000 |
| | Intercept | 2.293 | 0.019 | [2.257, 2.329] | 1.000 |
| a (boundary separation) | Reward (z) | 0.172 | 0.004 | [0.164, 0.181] | 1.000 |
| | Travel time (c) | 0.001 | 0.000 | [0.000, 0.002] | 1.000 |
| | Trait anxiety (z) | −0.006 | 0.003 | [−0.011, −0.001] | 1.000 |
| | Intercept | 0.955 | 0.009 | [0.938, 0.973] | 1.000 |
| t (non-decision time) | — | 0.216 | 0.001 | [0.213, 0.218] | 1.000 |
| z (starting point) | — | 0.527 | 0.004 | [0.520, 0.535] | 1.000 |

**Supplementary table.20: Model comparison table for HSSM in US population**

| **Parameters in model** | **p_LOO** | **LOO Metric (elpd_loo)** | **WAIC Metric** |
|---|---|---|---|
| Trait anxiety | −1035.32 | 8.19 | −1035.32 |
| Reward + Travel time | 6589.26 | 7.87 | 6589.26 |
| **Reward + Travel time + Trait anxiety** | **6591.89** | **9.50** | **6591.89** |
